# Diet and heat - one neuronal subset two responses

**DOI:** 10.1101/2020.11.18.388017

**Authors:** Elodie Prince, Jenny Kretzschmar, Laura C. Trautenberg, Marko Brankatschk

**Author notes:** iBV - Institut de Biologie Valrose, CNRS UMR7277, Inserm U1091, UNS - Bâtiment Centre de Biochimie, Faculté des sciences - Parc Valrose - 06108 Nice Cedex 2, France.

## Abstract

The Insulin signal cascade is one of the best studied metabolic circuits, and shows a remarkable high molecular and functional conservation across the animal kingdom. Insulin-producing cells respond directly to nutritional cues in circulation and receive modulatory input from connected neuronal networks. Neuronal control is rapid and integrates a wide range of variables including dietary change or environmental temperature. However, despite various detailed studies that demonstrated the potential of neuronal regulation the physiological relevance of this circuit remains elusive.

In *Drosophila*, Insulin-like peptide 7 (dIlp7)-producing neurons are wired with Insulin-producing cells. We found a dual role for this neuronal subset: a.) activated dilp7-producing neurons are required to facilitate development at high temperatures, and if confronted with calorie-rich food that represses neuronal activity b.) their product, dIlp7, regulates Insulin signalling levels. Our work shows that Insulin-producing cells not simply integrate signals from circulating nutritional cues and neuronal inputs, but switch to neuronal control in response to dietary composition.

## INTRODUCTION

Thriving in new ecological systems, *Drosophila* learned to utilize local food. Such resources vary in their composition in response to environmental temperature changes (1–3). Wild fruit flies feed on rotting fruits *i.e.* a composite diet of microbes and plants. As such food sources are erratic, adult fruit flies also feed on alternative carbohydrate sources, such as yeast-free pollen (4,5). Pollen represents a type of calorie-rich plant food characterized by unsaturated fatty acids and phytosterols (6,7). Previously, it has been shown that *Drosophila* struggle to survive on corresponding experimental diets at high temperatures (1) and that wild flies seek yeast-based food in the summer (3). Although global warming is the main problem our society is facing in this 21st century, its consequence on the animal kingdom is not understood (8,9). In this paper, we provide new insights in the dual response of one neuronal subset to diet and changes of environmental temperatures. With increasing temperatures, fruit flies accelerate their metabolism to acclimatize. By feeding on yeast, *Drosophila* can maintain high <u>I</u>nsulin <u>s</u>ignaling (IS) levels required to shuttle efficiently nutritional cues (such as sugars in circulation) into cells (1,10–12). *Drosophila* produces seven different <u>I</u>nsulin-<u>l</u>ike <u>p</u>eptides (dIlps) in response to dietary signals. The structure and function of dIlps, as well as the putative precursor protein, are similar to human Insulin and <u>I</u>nsulin-like <u>g</u>rowth <u>f</u>actor (IGF) peptides or pre-pro-Insulin, respectively; thus, implying an analogous organization of the final secreted peptide (12–14). Some dIlps play exclusive developmental roles, such as dIlp1 (15–17), dIlp 4 (12) and dIlp8 (18–21), and their production is restricted in time. Others, such as dIlp6 are associated with nutritional disorders and counteract the group of canonical dIlp peptides (22). Adult flies express four neuronal dIlps: dIlp2,3,5 and 7. DIlp2,3 and 5,produced by neuronal <u>I</u>nsulin-<u>p</u>roducing <u>c</u>ells (IPCs), are secreted into circulation (14,23) and resemble the functional role of the vertebrate Insulin, secreted by the pancreatic Beta cells (14). DIlp7 is produced by specialized <u>d</u>Ilp<u>7</u>-<u>n</u>eurons (D7Ns) and is the only dIlp peptide showing structural conservation across species (13,24,25). Some D7Ns are wired with IPCs whereas others terminate in the periphery (23,26–29). Therefore, dIlp7 may regulate IS at multiple levels, by directly controlling IPC activity or stimulating the IS in the periphery (23,27).

The intracellular IS cascade is conserved and consists of molecular components known from the vertebrate pathway (13,30). Activated <u>In</u>sulin <u>receptor</u> (InR) recruits its adaptor protein, Chico - the functional counterpart of vertebrate <u>I</u>nsulin <u>r</u>eceptor <u>s</u>ubstrate (IRS) proteins - which conveys the signal into either the Ras- or PI3K-dependent pathway (31). The Ras-dependent pathway promotes mitogenic metabolism characterized by catabolic cellular turnover rates (32). PI3K signaling promotes anabolic processes that buildup cellular energy stores (33).

Here, we investigate IS in the scope ofdifferent diets and changing environmental temperatures. We show that yeast products can compensate for the loss of dIlp2, 3 and 5 at moderate temperatures. However, to endure heat stress, all three dIlps and yeast are required. In addition, we show that high intracellular Calcium (Ca^2+^) levels in D7Ns are essential to develop at high temperatures. On yeast-free diets, D7Ns have low Ca^2+^ levels and in consequence, *Drosophila,* on such diets, do not develop.

We found that dIlp7 is essential to survive on yeast-free diet and identified the <u>l</u>eucin-rich repeat-containing <u>G</u> protein-coupled <u>r</u>eceptor <u>3</u> (Lgr3) as its potential target. We speculate that dIlp7/Lgr3 signalling is maintaining systemic IS levels by regulating dIlp2 and 5 activity. Taken together, we propose that D7Ns form a neurosecretory network essential to sustain *Drosophila* in absence of dietary yeast and changing environmental conditions.

## RESULTS

### Yeast food is not sufficient to replace IPC activity at high temperatures

*Drosophila* larvae kept on <u>Y</u>east <u>f</u>ood (YF) show high systemic <u>I</u>nsulin <u>s</u>ignalling (IS) levels, and can endure heat stress. It was shown that yeast lipids facilitate the secretion of *<u>D</u>rosophila* <u>I</u>nsuline-<u>l</u>ike <u>p</u>eptides (dIlps) but it remained unclear if high dIlp levels are essential to thrive at high temperatures (1). To test if <u>I</u>nsuin-<u>p</u>roducing <u>c</u>ell (IPC) activity is required for flies to survive in warm environments, we have silenced or hyper-activated IPCs by expressing either the inward rectifier potassium channel Kir2.1 (34) or the thermo-sensitive <u>t</u>ransient <u>r</u>eceptor <u>p</u>otential cation channel, TRPA1 (35,36). As expected, we found that on YF, at 28°C, IPC activity is essential for larval survival (Supplementary Figure S1). To investigate if IPC activity is required throughout larval development, we have induced IPC silencing at different larval stages. At high temperatures (28°C), IPC activity is essential for larval survival, whereas continuous high calcium (Ca^2+^) levels in these cells are of no consequence (Supplementary Figure S1). To investigate if IPC activity is required throughout larval development, we have induced IPC silencing at different larval stages. At 28°C, high Ca^2+^ levels in IPCs are essential for larval survival at any tested time point (Supplementary Figure S1). To evaluate the contribution of individual dIlps, we have tracked the development of different *ΔdIlp* mutants at comfortable temperatures (20°C) or upon heat stress (28°C). At 20°C, all tested mutants, including *ΔdIlp*^*2*^, *ΔdIlp*^*3*^, *ΔdIlp*^*5*^, *ΔdIlp*^*2-3*^ or *ΔdIlp*^*2-3,5*^, survive (at 20°C, n_larva_=132, control (*Cntr*)=132 ±0% and n_larva_= 140, *ΔdIlp*^*2-3,5*^=72.1 ±4.3%, p=****); although the developmental rate of *ΔdIlp*^*2-3,5*^ is much slower and produces small larvae and pupae with respect to controls (Figure 1 and Supplementary Figure S1 and (37,38).

**Figure 1.**
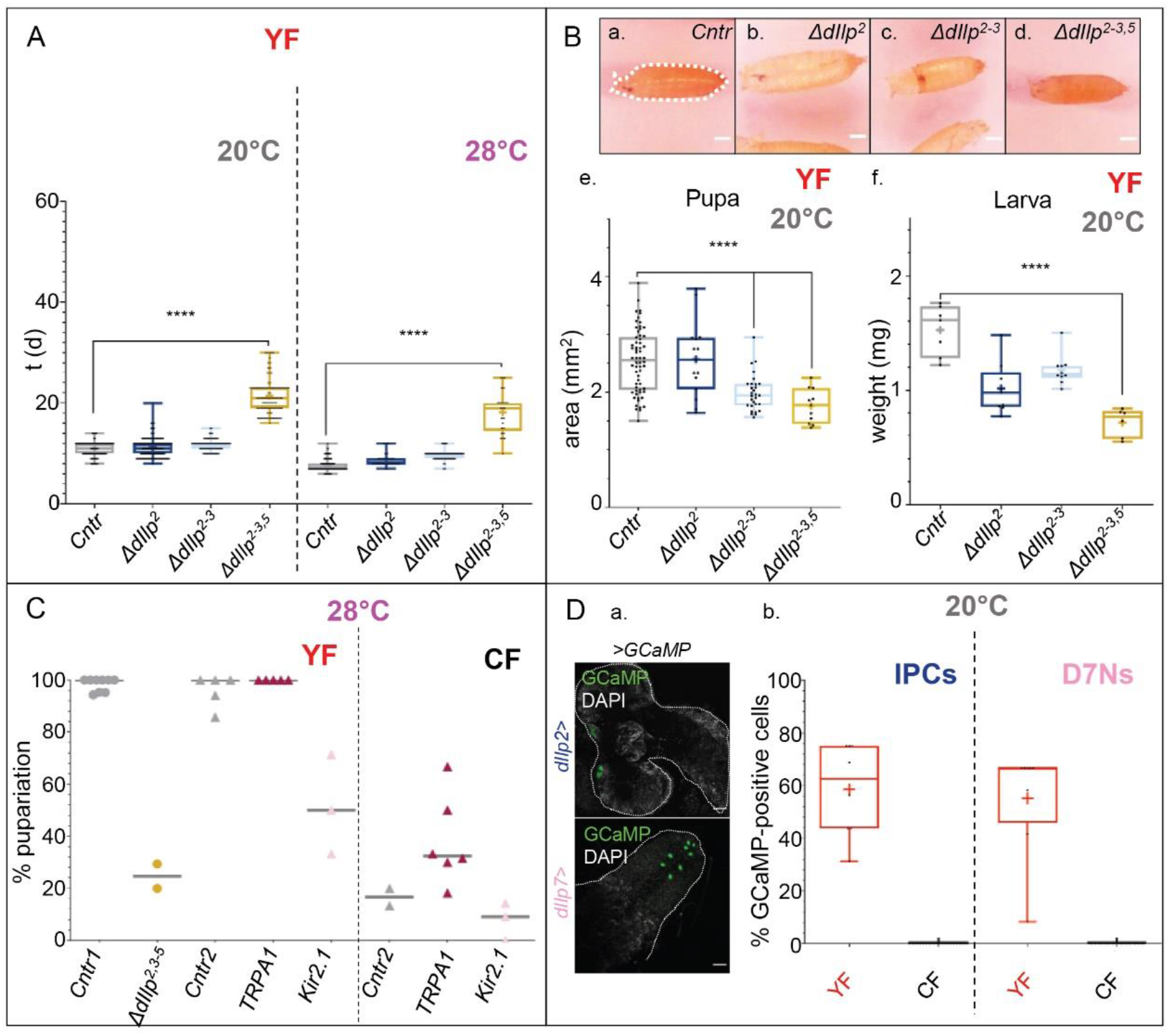
Activated D7Ns facilitate larval development at high temperatures. **(A).** Plotted is the developmental speed in days (d) to reach the pupal stage of FOXO^*mCherry*^ (*Cntr*, grey) and *ΔdIlp* mutant larvae grown on YF at 20 or 28°C. Each black dot represents one tracked larva. *Cntr* (grey; median^20°C^=11d; median^28°C^=7d), *ΔdIlp*^*2*^ (dark-blue; median^20°C^=11d; median^28°C^=8d), *ΔdIlp*^*2-3*^ (light-blue, median^20°C^=12d; median^28°C^=10d) and *ΔdIlp*^*2-3,5*^ (yellow; median^20°C^=21d; median^28°C^=19d). Statistics: Kruskal-Wallis test p<0.0001 ****; Dunn’s multiple comparison test: *Cntr vs ΔdIlp*^*2*^, p^20°C^>0.9999 ns, p^28°C^=0.0001 ***; *Cntr vs ΔdIlp*^*2-3*^, p^20°C^=0.0159 *, p^28°C^ 0.0001 ****; *Cntr vs ΔdIlp*^*2-3,5*^: p^20°C^, p^28°C^ <0.0001 ****. **(B). a.-d.** Shown are representative pictures of *FOXO*^*mCherry*^ (a, *Cntr*), *ΔdIlp*^*2*^ (b), *ΔdIlp*^*2-3*^ (c) and *ΔdIlp*^*2-3,5*^ pupae (d) raised on YF at 20°C. Scale bars=0.5mm. **e.** Plotted is the area of pupae grown on YF at 20°C. Each black dot represents one pupa. *Cntr* (grey; median=2.6), *ΔdIlp*^*2*^ (dark-blue; median=2.6), *ΔdIlp*^*2-3*^ (light-blue; median= 1.9) and *ΔdIlp*^*2-3,5*^ (yellow; median=1.8). Statistics: Kruskal-Wallis test p<0.0001 ****; Dunn’s multiple comparison test: *Cntr vs ΔdIlp*^*2*^, p>0.9999 ns; *Cntr vs ΔdIlp*^*2-3*^, p<0.0001 ****; *Cntr vs ΔdIlp*^*2-3,5*^, p<0.0001 ****. **f.** Plotted is the larva weight per larva calculated from n pool of larvae collected and grown on YF at 20°C. Each dot represents one pool. *Cntr* (grey; median=1.6), *ΔdIlp*^*2*^ (dark-blue; median=1,0), *ΔdIlp*^*2-3*^ (light-blue; median=1.1) and *ΔdIlp*^*2-3,5*^ (yellow; median=0.8). Statistics: Kruskal-Wallis test p<0.0001 ****; Dunn’s multiple comparison test: *Cntr vs ΔdIlp*^*2*^, p=0.0044 **; *Cntr vs ΔdIlp*^*2-3*^, p=0.1488 ns; *Cntr vs ΔdIlp*^*2-3,5*^, p<0.0001 ****. **(C).** Plotted is the pupariation rate (%) of larvae reared on YF or CF at 28°C. *FOXO*^*mCherry*^ (*Cntr1*, grey dots) and the *ΔdIlp*^*2-3,5*^ (yellow dots), *dIlp7-Gal4/+* (*Cntr2*, grey triangles), *dIlp7>>TRPA1* (*TRPA1*, dark-pink triangles) and *dIlp7>>Kir2.1,tub80ts* (*Kir2.1*, light-pink triangles). Each mark represents one experiment; the median is depicted as a grey bar. *Cntr1* (median=100%), *ΔdIlp*^*2-3,5*^ (median=24.7%), *Cntr2* (YF, median=100%; CF, median= 16.65%), *TRPA1* (YF, median=100%; CF, median=32.45%), *Kir2.1* (YF, median=50%; CF, median=9.10%). Statistics: Mann-Whitney test: *Cntr1 vs ΔdIlp*^*2-3,5*^, p=0.0182 *; Kruskal-Wallis test p=0.0031 **, Dunn’s multiple comparison test: *Cntr2 vs TRPA1*, p=0.7091 ns; *Cntr vs Kir2.1*, p=0.0549 ns (YF); Kruskal-Wallis test p=0.0069 **, Dunn’s multiple comparison test: *Cntr2 vs TRPA1*, p=0.3138 ns; *Cntr vs Kir2.1*, p=0.9484 ns (CF). **(D). a.** Representative Z-projection confocal image of GCaMP-positive cells (green) in brains from larvae expressing GCaMP in IPCs (top, *dIlp2>GCaMP*; stack=47.5μm) or D7N (bottom, *dIlp7>GCaMP*; stack=36.1μm). Larvae were kept on YF at 20°C. All samples were stained with DAPI (grey). Scale bars=25μm. **b.** Plotted is the percentage of GCaMP-positive IPCs or D7Ns in larvae reared on YF (red) or CF (black) at 20°C. IPCs (n_YF_=8; median=62.55%; n_CF_=23; median=0%); D7Ns (n_YF_=8; median=66.7%; n_CF_=20; median=0%). Statistics, Wilcoxon-test, YF *vs* CF, p=0.0078 ** (IPCs, D7Ns).

At 28°C, tested *ΔdIlp*^*2*^, *ΔdIlp*^*3*^ or *ΔdIlp*^*5*^ larvae, kept on YF, show no developmental defects (Supplementary Figure S1). However, we found that heat stress slows the developmental rate and reduces survival of *ΔdIlp*^*2-3*^, and that a majority of *ΔdIlp*^*2-3,5*^ animals perish (Figure 1).

Yeast-free food fed animals have limited IPC activity, low systemic IS levels and show decreased survival at high temperatures (1). To rule out the effects of yeast products, we have repeated all experiments using <u>C</u>orn <u>f</u>ood (CF), a yeast-free diet that resembles the ingredient proportions found in YF ((39); Table 1 and Table 2, respectively). At 28°C, controls are unable to survive on CF. Furthermore, induced expression of dIlp2 in IPCs is only in part promoting larval development (Supplementary Figure S1). Taken together, to mount an effective heat response, *Drosophila* relies on dietary yeast products and its complement of IPC dIlp peptides.

**Table 1.**
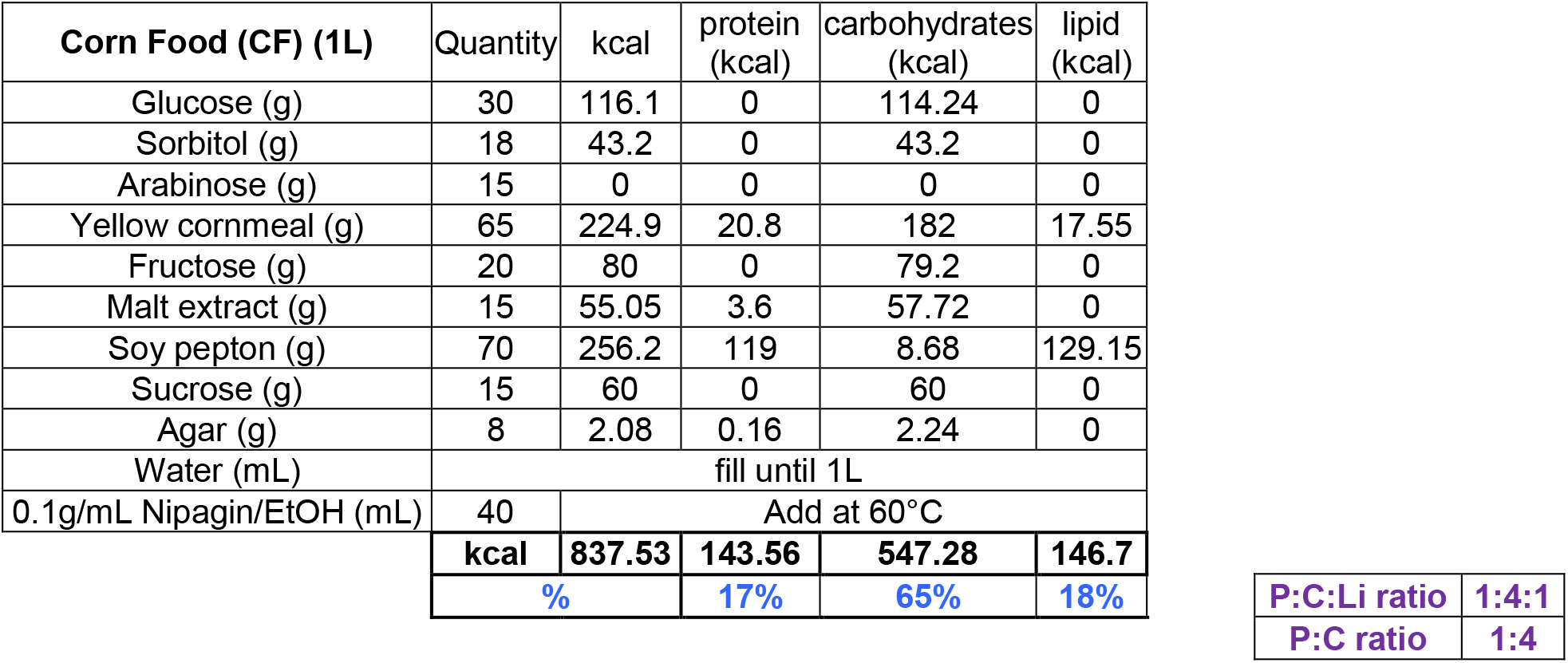
Corn food recipe. Recipe for 1 liter of corn food. The calories (in kcal) brought by each ingredient have been also calculated. At the bottom, in black bold, the totals (in kcal) are shown. In blue are indicated the proportion (%) of kcal of protein, carbohydrates or lipids on the total of amount calories. In purple are shown the protein: carbohydrate: lipid and protein: carbohydrate ratios. Abbreviations: EtOH, ethanol; P, proteins; C, carbohydrates; Li, lipids.

| **Corn Food (CF) (1L)** | Quantity | kcal | protein (kcal) | carbohydrates (kcal) | lipid (kcal) |
| --- | --- | --- | --- | --- | --- |
| Glucose (g) | 30 | 116.1 | 0 | 114.24 | 0 |
| Sorbitol (g) | 18 | 43.2 | 0 | 43.2 | 0 |
| Arabinose (g) | 15 | 0 | 0 | 0 | 0 |
| Yellow cornmeal (g) | 65 | 224.9 | 20.8 | 182 | 17.55 |
| Fructose (g) | 20 | 80 | 0 | 79.2 | 0 |
| Malt extract (g) | 15 | 55.05 | 3.6 | 57.72 | 0 |
| Soy pepton (g) | 70 | 256.2 | 119 | 8.68 | 129.15 |
| Sucrose (g) | 15 | 60 | 0 | 60 | 0 |
| Agar (g) | 8 | 2.08 | 0.16 | 2.24 | 0 |
| Water (mL) | fill until 1L |  |  |  |  |
| 0.1g/mL Nipagin/EtOH (mL) | 40 | Add at 60°C |  |  |  |
|  | **kcal** | **837.53** | **143.56** | **547.28** | **146.7** |
|  | **%** |  | **17%** | **65%** | **18%** |

**Table 1. Corn food recipe**
|  |  |
| --- | --- |
| **P:C:Li ratio** | **1:4:1** |
| **P:C ratio** | **1:4** |

**Table 2.**
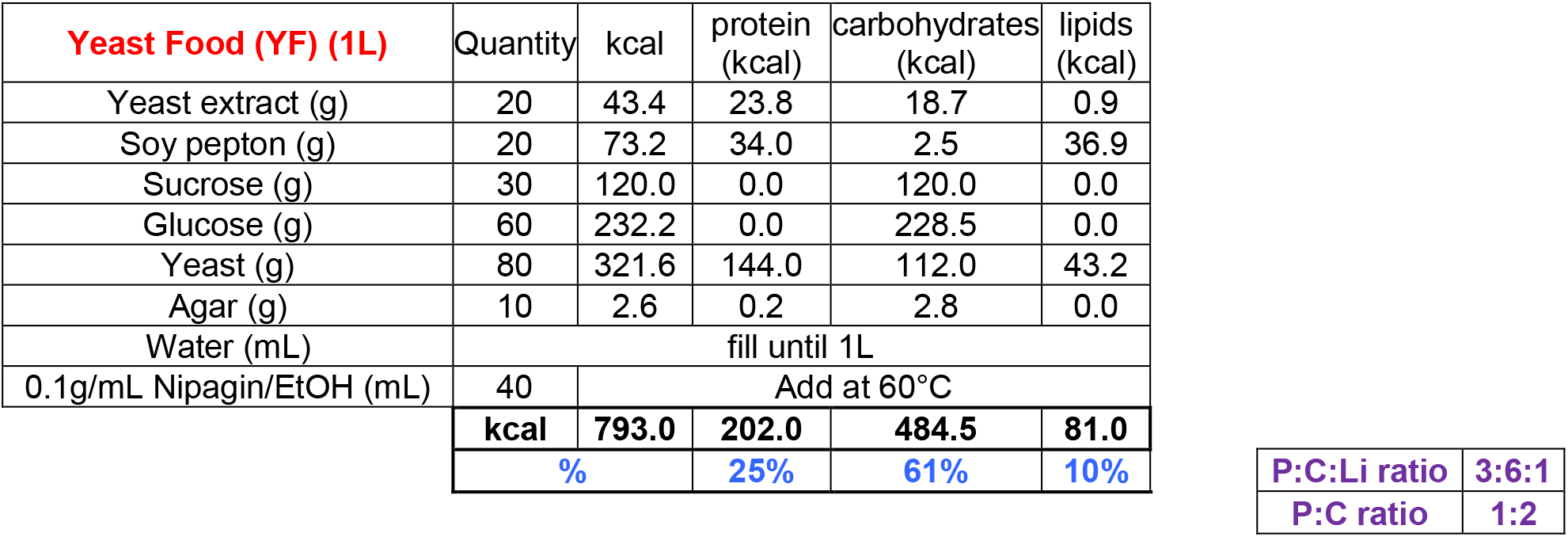
Yeast food recipe. Recipe for 1 liter of corn food. The calories (in kcal) brought by each ingredient have been also calculated. At the bottom, in black bold, the totals (in kcal) are shown. In blue are indicated the proportion (%) of kcal of protein, carbohydrates or lipids on the total of amount calories. In purple are shown the protein: carbohydrate: lipid and protein: carbohydrate ratios. Abbreviations: EtOH, ethanol; P, proteins; C, carbohydrates; Li, lipids.

| **Yeast Food (YF) (1L)** | Quantity | kcal | protein (kcal) | carbohydrates (kcal) | lipids (kcal) |
| --- | --- | --- | --- | --- | --- |
| Yeast extract (g) | 20 | 43.4 | 23.8 | 18.7 | 0.9 |
| Soy pepton (g) | 20 | 73.2 | 34.0 | 2.5 | 36.9 |
| Sucrose (g) | 30 | 120.0 | 0.0 | 120.0 | 0.0 |
| Glucose (g) | 60 | 232.2 | 0.0 | 228.5 | 0.0 |
| Yeast (g) | 80 | 321.6 | 144.0 | 112.0 | 43.2 |
| Agar (g) | 10 | 2.6 | 0.2 | 2.8 | 0.0 |
| Water (mL) | fill until 1L |  |  |  |  |
| 0.1g/mL Nipagin/EtOH (mL) | 40 | Add at 60°C |  |  |  |
|  | **kcal** | **793.0** | **202.0** | **484.5** | **81.0** |
|  | **%** |  | **25%** | **61%** | **10%** |

**Table 2. Yeast food recipe**
|  |  |
| --- | --- |
| **P:C:Li ratio** | **3:6:1** |
| **P:C ratio** | **1:2** |

### DIlp7-neurons are required to provide heat resistance in *Drosophila*

<u>D</u>Ilp<u>7</u>- producing <u>n</u>eurons (D7Ns) are wired with IPCs in larval and adult stages and have a modulative function (23,26,27). To test if D7N activity influences the resistance to high temperatures, we have silenced these cells and tracked larval development on YF at 28°C. We found that low D7N activity results in reduced larval survival (Figure 1). However, the developmental speed of surviving animals is comparable to controls (Supplementary Figure S1). *Drosophila* kept on yeast-free food show low IS levels and are sensitive to heat stress (1). Therefore, we speculated that D7N, on such yeast-free diets, mightbe inactive and that induced D7N activity could increase heat resistance. To test our idea, we expressed the fluorescent Ca^2+^ reporter, GCaMP, in D7Ns in larval brain tissues (10). We found that D7N never show high Ca^2+^ levels on CF contrary to flies kept on YF (Figure 1). To increase Ca^2+^ yields in D7Ns, we expressed the temperature-sensitive Ca^2+^ channel, TRPA1, and assayed the survival of larvae kept on yeast-free CF ((39) and Table recipes). Interestingly, survival rates of *dIlp*^*7*^>>*TRPA1* (*TRPA1*) larvae increased significantly (Figure 1). To investigate if dIlp7 is essential to provide heat resistance, we tracked *ΔdIlp*^*7*^ larvae at 20°C and 28°C. We found that *ΔdIlp*^*7*^ kept on YF show no phenotypes; however, *ΔdIlp*^*7*^ are unsuccessful on CF (Figure 2 and Supplementary Figure S2). We conclude that D7Ns are required to acclimatize at high temperatures. In addition, although not involved in temperature response, dIlp7 peptide produced by D7Ns is essential to promote survival on yeast-free diets.

**Figure 2.**
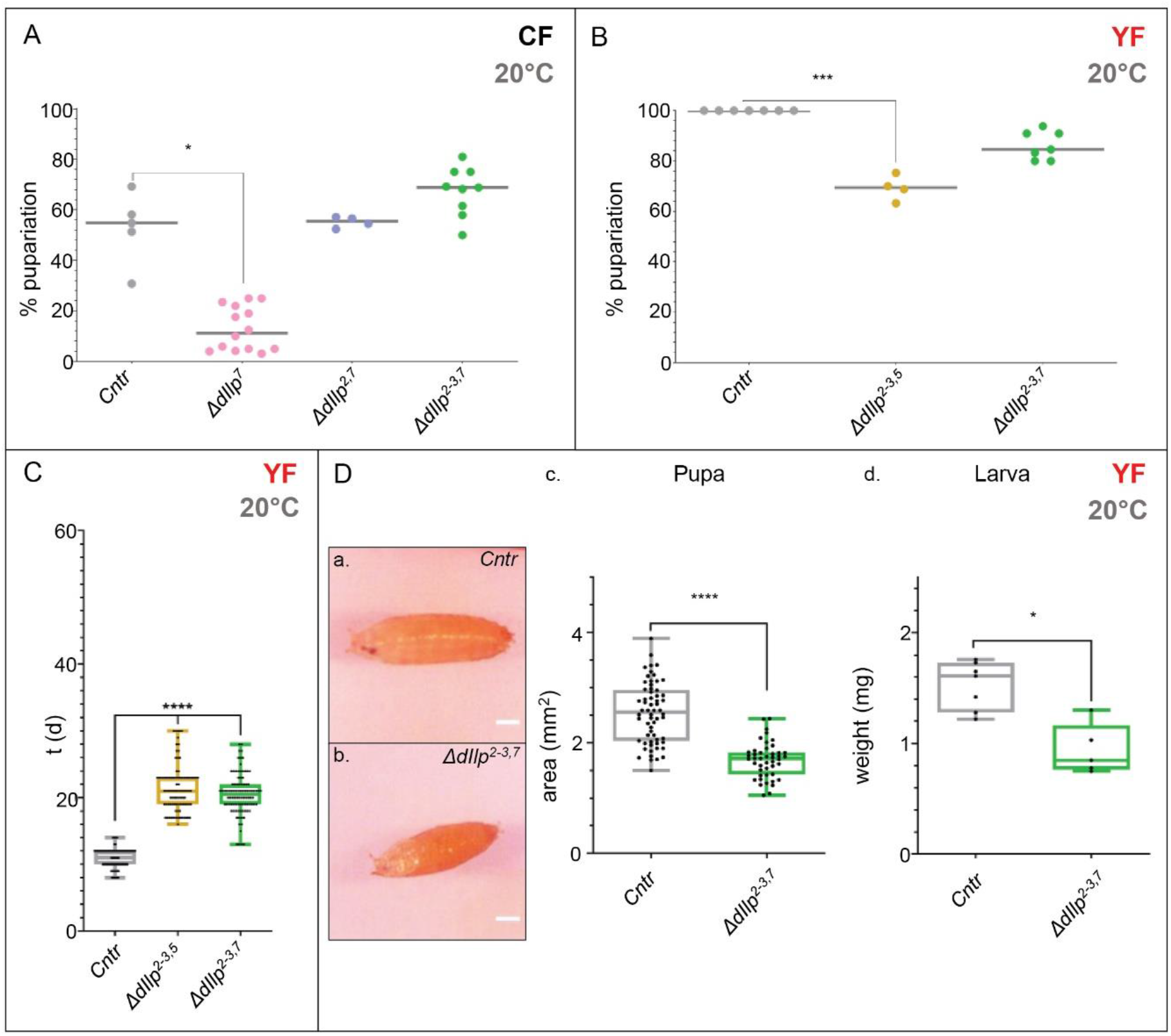
On CF dIlp7 peptide is instructive for IPCs. **(A).** Plotted is the pupariation rate in % (median=grey bar; each dot represents one experiment) of *FOXO*^*mCherry*^ (*Cntr*, grey), *ΔdIlp*^*7*^ (*pink*), *ΔdIlp*^*2,7*^ (blue) and *ΔdIlp*^*2-3,7*^ (green) larvae reared on CF at 20°C. *Cntr* (median=54.80%), *ΔdIlp*^*7*^ (median=11.25%), *ΔdIlp*^*2,7*^ (median=55.50%) and *ΔdIlp*^*2-3,7*^ (median=68.80%). Statistics: Kruskal-Wallis, p<0.0001 ****; Dunn’s multiple comparison test, *Cntr* vs *ΔdIlp*^*7*^: p=0.0183 *, *Cntr* vs *ΔdIlp*^*2,7*^: p>0.9999 ns, *Cntr* vs *ΔdIlp*^*2-3,7*^: p=0.8530 ns **(B).** Plotted are the pupariation rates in% (median=grey bar; each dot represents one experiment) of *FOXO*^*mCherry*^ (*Cntr*, grey), *ΔdIlp*^*2-3,5*^ (yellow) and *ΔdIlp*^*2-3,7*^ (green) larvae reared on YF at 20°C. *Cntr* (median=100%), *ΔdIlp*^*2-3,5*^ (median=69.40%) and *ΔdIlp*^*2-3,7*^ (median=84.60%). Statistics: Kruskal-Wallis, p<0.0001 ****; Dunn’s multiple comparison test, *Cntr* vs *ΔdIlp*^*2-3,5*^: p<0.0001 ****, *Cntr* vs *ΔdIlp*^*2-3,7*^: p<0.0001 ****. **(C).** Plotted is the developmental speed in days (d) to reach the pupal stage of *FOXO*^*mCherry*^ (*Cntr*, grey), *ΔdIlp*^*2-3,5*^ (yellow) and *ΔdIlp*^*2-3,7*^ (green) larvae reared on YF at 20°C. Each dot represents one tracked larva. *Cntr* (median= 11d), *ΔdIlp*^*2-3,5*^ (median=21d), *ΔdIlp*^*2-3,7*^ (median=20.5d). Statistics: Kruskal-Wallis, p<0.0001 ****; Dunn’s multiple comparison test, *Cntr* vs *ΔdIlp*^*2-3,5*^: p<0.0001 ****, *Cntr* vs *ΔdIlp*^*2-3,7*^: p<0.0001 ****. **(D). a-b.** Representative pictures of *FOXO*^*mCherry*^ (a, Control, *Cntr*) and *ΔdIlp*^*2-3,7*^ (b) larvae reared on YF at 20°C. Scale bars=0.5mm. **c.** Area of *FOXO*^*mCherry*^ (*Control*, *Cntr*, grey) and *ΔdIlp*^*2-3,7*^ (green) larvae reared on YF at 20°C. Each dot represents one experiment. The area of pupae, grown on YF at 20°C, has been measured as depicted in Fig. 1Ba, dashed line. *Cntr* (median=2.6) and *ΔdIlp*^*2-3,7*^ (median=1.7). Statistics: Mann-Whitney test, p<0.0001 ****. **d.** Plotted is the larva weight per larva calculated from n pool of larvae collected and grown on YF at 20°C. Each dot represents one pool. *Cntr* (grey; median=1.6) and *ΔdIlp*^*2-3,7*^ (green; median=0.850). Statistics: Mann-Whitney test, p=0.0101 *.

### On yeast-free food, dIlp2 and dIlp7 are negative regulators of dIlp5

To investigate the role of dIlps in animals feeding on yeast-free diets, we performed a metabolic screen of *ΔdIlp* mutants on CF at 20°C. We found that, in addition to *ΔdIlp*^*7*^, three other mutants struggle to thrive on CF, including Δ*dIlp*^*2*^, Δ*dIlp*^*3*^ and Δ*dIlp*^*5*^ (Figure 2). To evaluate potential genetic interactions between dIlp7 and each of these candidates, we have created *ΔdIlp*^*2,7*^, *ΔdIlp*^*3,7*^ and *ΔdIlp*^*5,7*^ double mutants. Astonishingly, the absence of dIlp2 and dIlp7 restored survival rates close to numbers shown by controls. In contrast, *ΔdIlp*^*3,7*^ and *ΔdIlp*^*5,7*^ animals did not perform better and *ΔdIlp*^*2,3*^ were unable to develop on CF (Figure 2; Supplementary Figures S1 and S2). To test, if the loss of dIlp2 and dIlp7 hyper-activates dIlp3 or dIlp5, we assayed the development of *ΔdIlp*^*2-3,5*^ and *ΔdIlp*^*2-3,7*^ triple mutants. In contrast to *ΔdIlp*^*2-3,5*^, *ΔdIlp*^*2-3,7*^ were able to survive on CF alike *ΔdIlp*^*2,7*^ or control larvae (Figure 2).

The presence of dIlp7 is required for survival on CF but not for individuals kept on YF. Hence, we tested the idea that yeast products compensate for the regulative role of dIlp7. To do so, we reared all created *ΔdIlp* mutants onto YF. We found that at 20°C, *ΔdIlp*^*5,7*^ are less successful, but did not reduce their developmental speed (Supplementary Figure S2). The lethality of *ΔdIlp*^*2-3,5*^ is comparable to *ΔdIlp*^*5,7*^; however, these triple mutants develop slower (Figure 1; Supplementary Figures S1 and S2). Although delayed in development, the survival of *ΔdIlp*^*2-3,7*^ is not affected (Figure 2). Pupae of both triple mutants are small in size (Supplementary Figure S1).

In conclusion, the data from our work indicate that on CF, dIlp5 can compensate for the loss of all other neuronal dIlps expressed in larval stages. Our findings crystallize dIlp2 and dIlp7 as negative regulators of dIlp5.

### DIlp7 is instructive for larval and adult feeding behavior on CF

Feeding behavior and intestinal food transit are two connected variables that can modulate the absorption efficiency of food compounds. It was shown that *Drosophila*, with genetically silenced D7Ns, change their feeding behavior with respect to controls (27). To test the feeding behavior of *ΔdIlp* mutants, we have measured larval mouth-hook contractions (40), and the appearance or disappearance of ingested color in the adult intestinal tracts (27). Transferred on CF, mouth-hook contractions of all tested *dIlp* mutants appeared slightly faster with respect to controls. Nevertheless, we found that ingestion of stained food in *ΔdIlp*^*7*^ mutant larvae takes longer compared to controls. (Supplementary Figure S2). To test if *ΔdIlp*^*7*^ adults show the same feeding behavior on CF, we recorded how fast colored food is consumed. *ΔdIlp*^*7*^ flies feed faster, but need more time to excrete material compared to controls (Figure 3). Interestingly, we have discovered a similar behavior for *ΔdIlp*^*2*^ mutants feeding on CF (Supplementary Figure S2). Taken together, these data show that dIlp7 and dIlp2 are important for larva and adult feeding behavior on yeast-free diet.

**Figure 3.**
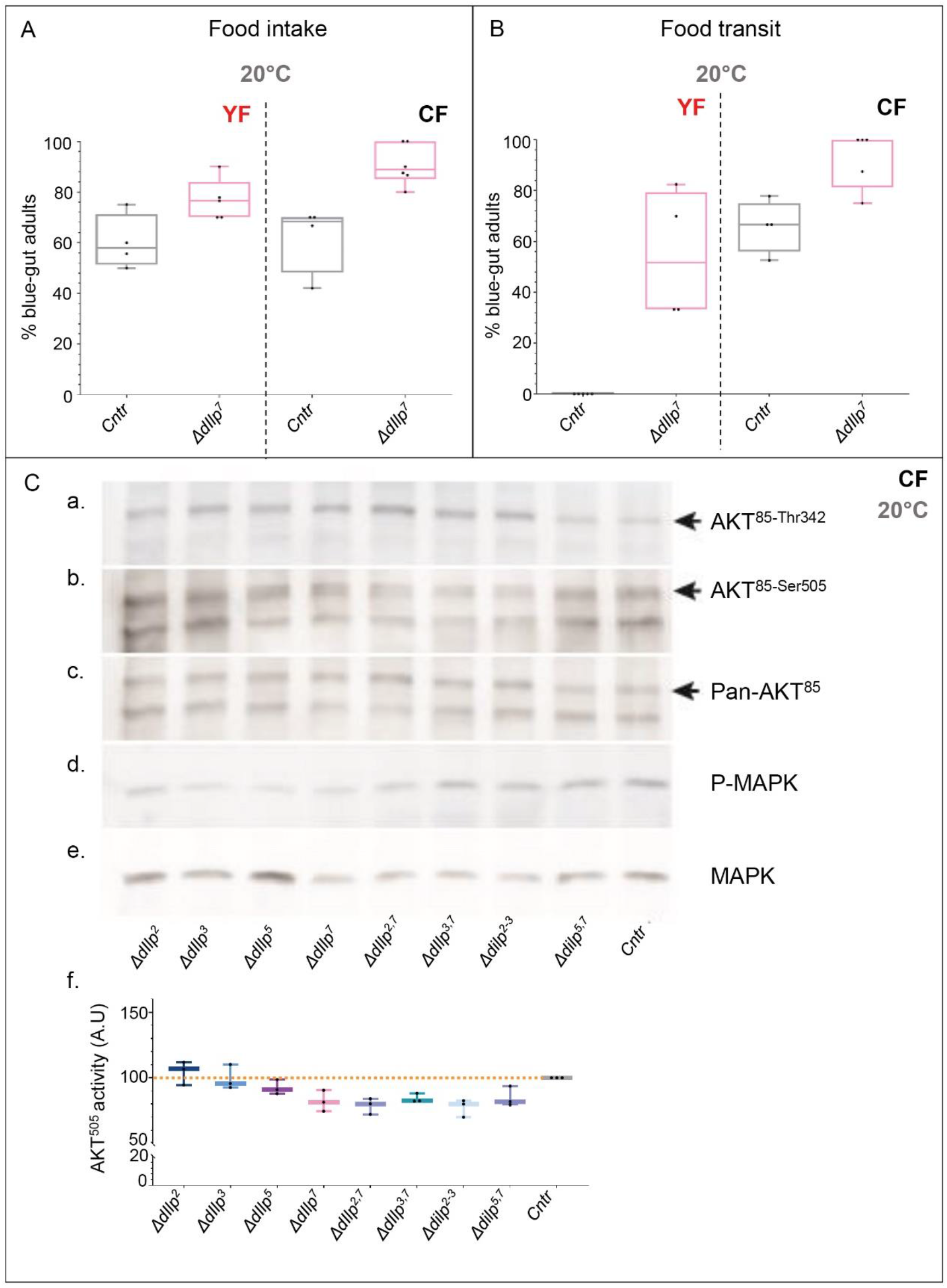
DIlp7 peptide regulates systemic IS on CF. **(A).** Plotted is the percentage of *CantonS* (grey, *Cntr*) and *ΔdIlp*^*7*^ (pink) adult females with blue guts, 4 hours after their transfer from uncoloured starvation plates to blue-stained YF (red, left) or CF (black, right). Each black dot = one experiment. *Cntr* (YF, median=57.80%; CF, median=68.35%) and *ΔdIlp*^*7*^ (YF, median=76.50%; CF, median=88.75%). Statistics: Mann-Whitney test, *Cntr vs ΔdIlp*^*7*^: pYF=0.0556 ns, p_CF_ =0.0095 **. **(B).** Plotted is the percentage of *CantonS* (grey, *Cntr*) and *ΔdIlp*^*7*^ (pink) adult females with blue guts 4 hours after their transfer from blue-stained YF (red, left) or CF (black, right) to uncoloured starvation plates. Each black dot = one experiment. *Cntr* (YF, median=0%; CF, median=66.70%) and *ΔdIlp*^*7*^ (YF, median=51.65%; CF, median=100%). Statistics: Mann-Whitney test, *Cntr vs ΔdIlp*^*7*^: p_YF_=0.0079 **, p_CF_=0.0317 *. **(C). a-e**. The phosphorylation states of AKT^85^ (a-c; marked by arrow) and MAPK (d, e) in *Control* (*Cntr*, *FOXO*^*mCherry*^) and *ΔdIlp* mutants kept on CF at 20°C were analysed by western blotting. Shown is the phosphorylation at the Thr^342^ (a, AKT^85-Thr342^), the Ser^505^ (b, AKT^85-Ser505^) site and total amount of AKT^85^ (c, Pan-AKT). In addition, phosphorylation (d.) and total yields (e.) of MAPK were detected. . **f.** The activity at the AKT^85-Ser505^ was quantified (in arbitrary units, A.U.) from 3 independent blots; each black dot=one experiment. Plotted is the relative activity of each genotype normalized to the control *FOXO*^*mCherry*^ (*Cntr*, grey, median=100%). *ΔdIlp*^*2*^ (dark-blue, median=106.7%), *ΔdIlp*^*3*^ (blue, median=95.60%), *ΔdIlp*^*5*^ (dark-purple, median=91.10%), *ΔdIlp*^*7*^ (pink, median=81.40%), *ΔdIlp*^*2,7*^ (blue, median=80.10%), *ΔdIlp*^*3,7*^ (green-blue, median=82.40%), *ΔdIlp*^*2-3*^ (light-blue, median=79.90%), *ΔdIlp*^*5,7*^ (light-purple, median=81.80%). Statistics: Kruskal-Wallis test, p=0.0120 *; Dunn’s multiple comparison test, *Cntr* vs *ΔdIlp*^*2*^, *ΔdIlp*^*3*^, *ΔdIlp*^*5*^: p>0.9999 ns, *Cntr* vs *ΔdIlp*^*7*^: p=0.1883 ns, *Cntr* vs *ΔdIlp*^*2,7*^: p=0.0934 ns, *Cntr* vs *ΔdIlp*^*3,7*^: p=0.5113 ns, *Cntr* vs *ΔdIlp*^*2-3*^: p=0.0596 ns, *Cntr* vs *ΔdIlp*^*5,7*^: p=0.3579 ns.

### DIlp7 and not dIlp2 regulates systemic Insulin signaling on CF

Food transition time might be critical for efficient absorption of nutrients, therefore we predicted that *ΔdIlp*^*2*^ and *ΔdIlp*^*7*^ mutants will show similar activity of metablic circuits such as the IS cascade. The phosphorylation state of AKT and MAPK allows to quantify the activity of anabolic and mitogenic IS activity, respectively. Therefore, we probed the phosphorylation state of AKT^85^ and MAPK in adults, fed on CF, by western-blotting. We found that the AKT^85-Thr342^ phosphorylation state remains unchanged in all tested genotypes (Supplementary Figure S3). In contrast, we detected reduced phosphorylation at the AKT^85-Ser505^ site of all tested *ΔdIlp*^*7*^ mutant genotypes and *ΔdIlp*^*2-3*^ mutants, while the AKT^85^ activity of *ΔdIlp*^*2*^, *ΔdIlp*^*3*^ and *ΔdIlp*^*5*^ mutants remains unchanged (Figure 3). In addition, we detected reduced MAPK phosphorylation for all *ΔdIlp* loss-of-function genotypes (Supplementary Figure S3). For comparison we repeated experiments performed on CF with YF to compare the activity of AKT^85^ and MAPK. We did not detect any meaningful phenotypes in most of *ΔdIlp* mutant regarding larval development and adult *ΔdIlp* mutants thrive on YF like controls (Supplementary Figure S3). Thus, we expected that neither *ΔdIlp* mutant shows differences in the activity of either enzyme compared to controls. We found that all tested genotypes kept on YF show similar AKT or MAPK phosphorylation levels (Supplementary Figure S3). Taken together, we conclude that YF compensates for the loss of individual dIlps. In contrast, on yeast-free diet (CF), adult flies rely on dIlp7 to adjust their feeding behavior and to maintain basic AKT and MAPK activity required to survive (data not shown).

### Lgr3 represents a potential target for dIlp7 on Insulin-producing cells

We speculated that on yeast-free food, dIlp7 regulates IPC activity by activating one of its three predicted candidate receptors: the *<u>D</u>rosophila* <u>I</u>nsulin-like <u>r</u>eceptor (dInR) (12), or one of the two predicted <u>G</u>-<u>p</u>rotein <u>c</u>oupled receptors (GPCR), <u>L</u>eucine-rich repeat-containing <u>G</u> protein-coupled <u>r</u>eceptor <u>3</u> (Lgr3) and <u>4</u> (Lgr4) (41,42). We decided to targeteach receptor in IPCs, by RNA interference (RNAi) using published validated genetic tools (43). The reduction of the dInR (InR^KD^) or Lgr4 (Lgr4^KD^) did not produce any phenotypes. InR^KD^ and Lgr4^KD^ larvae developed as fast as controls; however, InR^KD^ individuals show a slight decrease in survival on both YF and CF compared to controls (Supplementary Figure S4). In contrast, we found that the loss of Lgr3 (Lgr3^KD^) slowed the development on YF and decreased pupariation success on CF (Figure 4).

**Figure 4.**
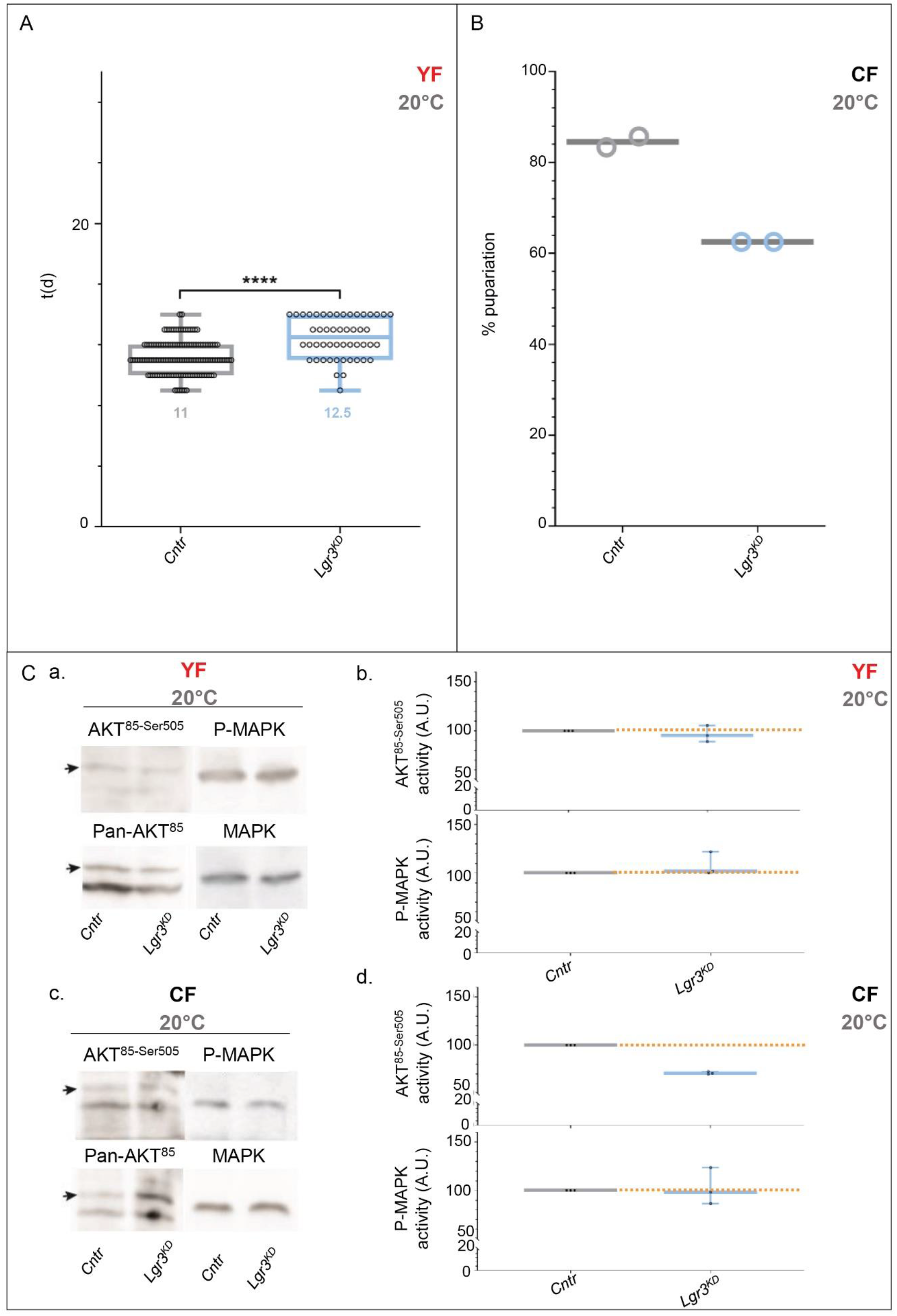
DIlp7 targets Lgr3 expressed by IPCs. **(A, B).** Plotted is the developmental speed in days (d, A) and the pupariation rate (%, B) of control *dIlp2-Gal4/+* (*Cntr*, grey) and *dIlp2>>Lgr3RNAi* (*Lgr3*^*KD*^, light-blue) larvae reared on YF (A) or on CF (B) at 20°C. **A.** Each black dot = one larva, the median is indicated. *Cntr* (median=11d) and *Lgr3*^*KD*^ (median=12.5d). Statistics: Mann-Whitney test: p<0.0001****. **B.** Each dot = one experiment. The median is indicated. *Cntr* (grey, median=84.5%) and *Lgr3*^*KD*^ (light-blue median=62.5%). **(C).** Shown are blots probed for AKT (a.) and MAPK (c.) of *Control* (*Cntr*, *FOXO*^*mCherry*^) and *dIlp2>>Lgr3RNAi (Lgr3*^*KD*^) flies. Flies were raised on YF (a, b) or CF (c, d) at 20°C. The phosphorylation at the AKT^85-Ser505^ and P-MapK site was quantified and relative activity plotted (**b,d).** each black dot = one experiment. Statistics: Mann-Whitney test, *Cntr* vs *Lgr3*^*KD*^, AKT^85-Ser505^: p=0.1000 ns; P-MAPK: p=0.7000 ns. On CF (d), *Lgr3*^*KD*^ (n-w=3; AKT^85-Ser505^, median=70.80%; P-MAPK, median=98.20%). Statistics, Mann-Whitney test, *Cntr* vs *Lgr3*^*KD*^, AKT^85-Ser505^: p=0.7000 ns; P-MAPK: p=0.1000 ns.

One accessible marker for stored cellular energy is the <u>T</u>ri<u>a</u>cyl<u>g</u>lyceride (TAG) level stored in lipid droplets. High IS levels correlate with high TAG yields. To test the idea that loss of Lgr3 reduces IS and hence, produces fewer TAGs, we have analysed lipid profiles of adult flies kept on YF. We found, that Lgr3^KD^ store less lipids with respect to controls (Supplementary Figure S4) indicating lower anabolic turnover rates. To investigate if Lgr3 modulates systemic IS, we probed the phosphorylation of AKT^85^ and MAPK in Lgr3^KD^ flies. We found that AKT^85^ phosphorylation of Lgr3^KD^ adults kept on YF at 20°C remained unchanged (Figure 4). On CF, the AKT^85-Ser505^ phosphorylation was reduced to levels assimilable to tested *ΔdIlp*^*7*^ mutant backgrounds (Figures 3 and 4). However, Lgr3^KD^ MAPK activity remained comparable to controls (Figure 4) indicating that dIlp7 should regulate MAPK by other means than Lgr3 expressed by the IPCs. Although of indirect evidence, our data lead us to the conclusion that on yeast-free diets, systemic IS is regulated by Lgr3 expressed in the IPCs.

## DISCUSSION

*Drosophila* expresses four neuronal <u>I</u>nsulin-<u>l</u>ike <u>p</u>eptides (dIlps) and we analyzed here the physiological role of dIlp7 produced by the dIlp7-neurons (D7Ns). We found that active D7Ns facilitate Insulin signalling (IS) and therefore, represent one component of a high-temperature-sensitive neuronal network similar to already described regulative circuits (44–46). Nonetheless, *ΔdIlp*^*7*^ mutants thrive at high temperatures similar to respective controls confirming that dIlp7 peptide is not crucial to withstand thermal treatment. We found that dIlp7 is essential to modulate IS in flies feeding on yeast-free diet, a food that allows only for low D7N activity and changes flies heat sensitivity.

Adult flies switch their feeding behavior in response to temperature and prefer yeast-free plant food during periods of cold temperatures (1,3). Moreover, *Drosophila* kept on plant food are unable to endure heat stress. We show that on <u>c</u>orn <u>f</u>ood (CF), D7Ns have low calcium (Ca^2+^) levels and induced activity of these cells enables more animals to survive heat. Interestingly, we found high dIlp7 levels in D7Ns regardless of the dietary input (Supplementary Figure S5). Hence, we wondered if dIlp7 plays any role for *Drosophila* feeding on yeast-free food. We show that the developmental success of *Δdilp*^*7*^ is strongly reduced on CF with respect to controls. It was shown that dIlp7 expression is strongly dependent on nutritional variables including protein:sugar ratio and caloric load (47). CF is rich in calories, its protein:sugar ratio is optimized and contains about 10% lipids ((39) and Table 1). Nevertheless, on CF, systemic Insulin levels are low and wild-type larvae struggle to complete their development (39,48). Astonishingly, we found that the absence of both dIlp2 and dIlp7 improved larval survival on CF up to numbers shown by controls. DIlp2 and dIlp5 have been described as highly redundant and target identical dIlp binding proteins (13). Nevertheless, the survival rate of *ΔdIlp*^*5,7*^ mutants kept on CF is low; thus, implicating that dIlp5 plays a different biological role than dIlp2 (49,50). In addition, *ΔdIlp*^*2-3,7*^ mutants are as successful as *ΔdIlp*^*2,7*^; thus, ruling out dIlp3 as a determining factor. Moreover, our results position dIlp5 as sufficient to rescue development since *ΔdIlp*^*2-3,5*^ and *ΔdIlp*^*2-3*^ hardly grow on CF. Taken together, on yeast-free food, larval development is controlled by dIlp7, which may regulate the activity of dIlp2 and 5.

To better understand how possibly dIlp7 regulates the activity of the IPC-produced dIlp2 and dIlp5, we have investigated three predicted targets of dIlp7 including the *<u>D</u>rosophila* <u>I</u>nsulin-like <u>r</u>eceptor (dInR) and the two <u>G</u> <u>p</u>rotein-<u>c</u>oupled <u>r</u>ectors GPCRs, Lgr3 and 4 (12,41,42). We found that the knock-down of Lgr3 in IPCs results in lower larval survival on CF. It is shown that Lgr3 is targeted by dIlp8 and plays an important role in larval development (18,21). However, *ΔdIlp*^*8*^ mutants performed as controls on CF (Supplementary Material/Excel sheet/dIlp mutants_Survival_CF20) and AKT^85-Ser505^ phosphorylation of adult Lgr3^KD^ is reduced to numbers shown by all tested *ΔdIlp*^*7*^ background mutants. On the other hand, measured MAPK activity of Lgr3^KD^ is comparable to controls. This result is contrasting findings made with *ΔdIlp*^*7*^ mutants, and we propose that secreted dIlp7 enters circulation (23,26) and regulates the mitogenic branch of the IS cascade. Nevertheless, future studies are required to demonstrate direct dIlp7/Lgr3 interaction at the molecular level.

Neuronal innervation of IPCs is established in many animals and modulates metabolic signals (23,45,51,52). Our findings indicate that food products can overwrite neuronal stimulation of IPCs. In *Drosophila*, we found a dual role for D7Ns: i.) these neurons facilitate the heat response of flies feeding on yeast and ii.) they form a metabolic circuit that enables adult flies to thrive on yeast-free diets if required. In mice and humans, pancreatic islets are directly innervated (51,52); however, the role of this neuronal stimulation in response to dietary cues is not understood. We have identified D7Ns and their product, dIlp7, important to regulate IS independently from the provided high caloric load and abundance in dietary sugars or proteins. Our findings provide new insights into the neuronal stimulation of Insulin-producing cells within a given ecological context and open the perspective to screen for nutritional cues capable to convey dietay signals to the brain.

## MATERIAS AND METHODS

### Stocks

If not stated, stocks were raised on normal food at 20-22°C (Table 3). The following lines were used: *OregonR* (#5), *w;foxo[mCherry]* (#80565) were provided by Bloomington Drosophila Stock Center (BDRC). Isogenized *Canton S* were shared by J.C. Billeter. *w*^*1118*^ and *yw* were provided by S. Eaton’s laboratory.

**Table 3.**
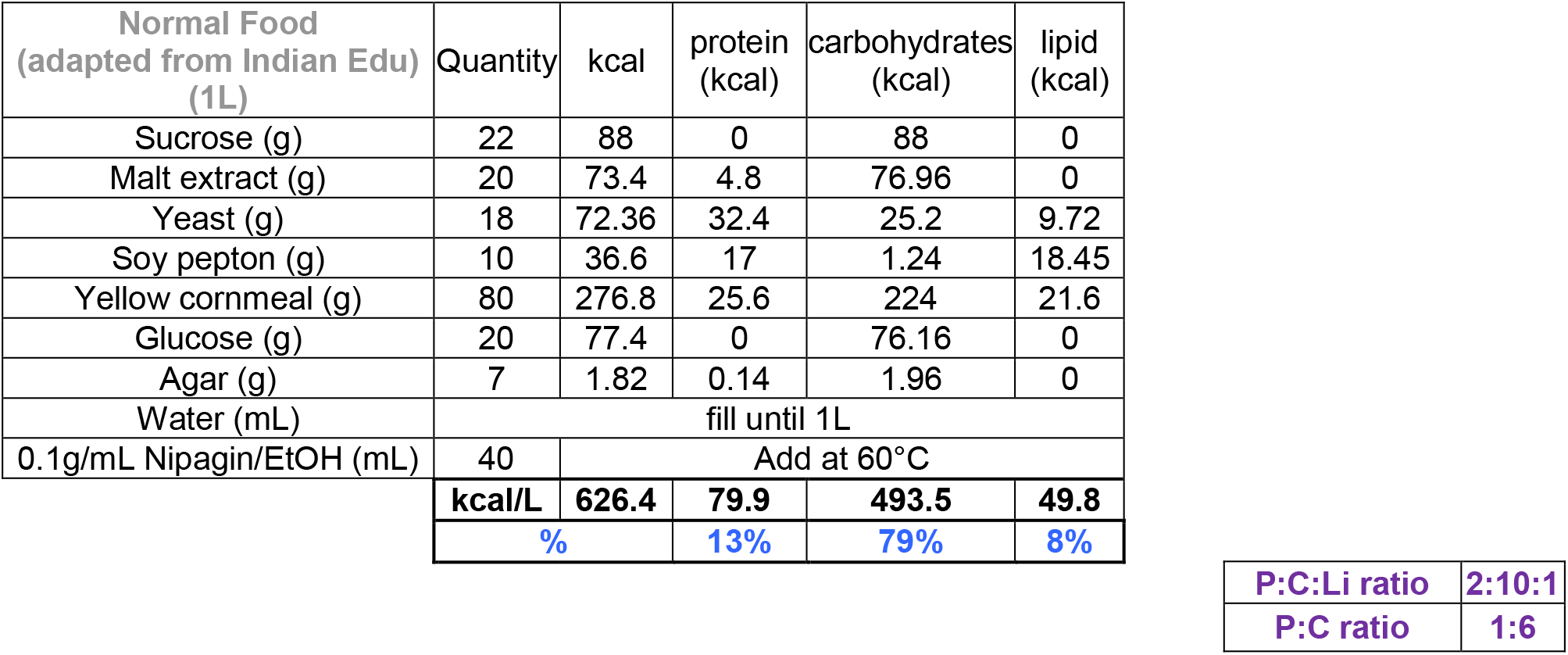
Normal food recipe. Recipe for 1 liter yeast food. Quantities of each ingredient (in g or mL) are indicated. The calories (in kcal) brought by each ingredient have been also calculated. At the bottom, in black bold, the totals (in kcal) are shown. In blue are indicated the proportion (%) of kcal of protein, carbohydrates or lipids on the total of amount calories. In purple are shown the protein: carbohydrate: lipid and protein: carbohydrate ratios. Abbreviations: EtOH, ethanol; P, proteins; C, carbohydrates; Li, lipids

| **Normal Food (adapted from Indian Edu) (1L)** | Quantity | kcal | protein (kcal) | carbohydrates (kcal) | lipid (kcal) |
| --- | --- | --- | --- | --- | --- |
| Sucrose (g) | 22 | 88 | 0 | 88 | 0 |
| Malt extract (g) | 20 | 73.4 | 4.8 | 76.96 | 0 |
| Yeast (g) | 18 | 72.36 | 32.4 | 25.2 | 9.72 |
| Soy pepton (g) | 10 | 36.6 | 17 | 1.24 | 18.45 |
| Yellow cornmeal (g) | 80 | 276.8 | 25.6 | 224 | 21.6 |
| Glucose (g) | 20 | 77.4 | 0 | 76.16 | 0 |
| Agar (g) | 7 | 1.82 | 0.14 | 1.96 | 0 |
| Water (mL) | fill until 1L |  |  |  |  |
| 0.1g/mL Nipagin/EtOH (mL) | 40 | Add at 60°C |  |  |  |
|  | **kcal/L** | **626.4** | **79.9** | **493.5** | **49.8** |
|  | **%** |  | **13%** | **79%** | **8%** |

**Table 3. Normal food recipe**
|  |  |
| --- | --- |
| **P:C:Li ratio** | **2:10:1** |
| **P:C ratio** | **1:6** |

*Drosophila* Insulin-like peptide mutants: *Δdilp*^*1*^-*Δdilp*^*8*^, *Δdilp*^*2-3*^ and *Δdilp*^*2-3,7*^ ((13); from S. Grönke); *Δdilp*^*2-3,5*^ from R. Kühnlein.

Gal4 lines: *w;dilp2-Gal4* ((14); #37516 from BDRC); *w;dilp7-Gal4* ((27); gift from I. Miguel-Aliaga); *w;rab3-Gal4* and w;*moody-Gal4* (from S.Eaton); *yw;actin-Gal4*(#4414, from BDRC) UAS-lines: *w;UAS-TrpA1* (#26264, from BDRC); *w;UAS-Kir2.1-egfp,tub80ts* (from S.Eaton); *yw;UAS-Dilp2* ((11); #80936 from BDRC); *w;UAS-GcAMP* (#32116 from BDRC); *yw;UAS-Dilp3* and *yw;UAS-Dilp5* (from S.Eaton); *w;UAS-Dilp7* (gift from I. Miguel-Aliaga); *yw;UAS-IR*^*DN*^ (#8253, from S.Eaton); *w;UAS-Lgr4RNAis* (v102681 and v108915; Vienna Drosophila Resource Center (VDRC)); *w;UAS-Lgr3shRNA* (v330603; from VDRC).

### Diets

Normal food was adapted from https://bdsc.indiana.edu/information/recipes/bloomfood.htm.l. (Table 3)

Yeast food was prepared as published by Brankatschk et al., 2018 (1) (Table 2).

Corn food recipe is described in Table 1 and in Trautenberg et al. 2020 (39).

### Development tracking experiments

Embryos were collected overnight on apple-juice agar plates (25% apple-juice, 1.5% agar, 0.4% p-Hydroxybenzoic acid methyl ester (nipagin)), bleached in 80% bleach/water for 30 seconds, rinsed and kept for 24 hours on starvation plates (25% apple juice, 10% Glucose, 1.5% agar, 0.4% nipagin) at 20-22°C. First-instar larvae were pipetted in 10μL 0.00005% Triton-X100/PBS solution and transferred in 96 well-plates containing food. One larva per well (containing 200μL of food) was deposited. Later, the plates were sealed with a poked ventilated plastic film and put in incubators set at the temperature of interest. Both developmental speed/rate (how many days each larva took to reach the pupal stage) and success (how many larvae succeed to reach the pupal stage on the total number of larvae plated on one plate).

Induction experiments: The egg progeny of crosses between *UAS-Kir2.1* or *Canton S* and *dilp2-Gal4*, *dilp7-Gal4* or *Canton S* were collected. Larvae and plates were prepared as described above. Plates were kept at 20°C for 24, 72 or 96 hours and then transferred to the 28°C-incubator.

### Statistics

Developmental success: For 20°C or 28°C/induction experiments, only experiments with minimum ten or six larvae tracked, respectively, were taken into account in the pool of data. Experiments out of the range (*i.e.* Mean + or − Standard deviation) were excluded of the normalized data. Mann-Whitney or Kruskal-Wallis tests were performed as statistical tests (PRISM Graphpad software).

Developmental speed: only speed data from experiments taken into account in the normalized data of the survival rates were picked. Mann-Whitney or Kruskal-Wallis tests were performed as statistical tests (PRISM Graphpad software).

### Immunohistochemistry

The larvae were dissected, turned as socks in fix solution and kept in fix solution for 20 minutes. The carcasses were stained by HrpCy5 (HRP-S5-1; NANOCS) or with DAPI (mounted in Vectashiel medium containing DAPI, vectorlab). The central nervous systems were dissected from the carcasses and mounted in 50% Glycerol/PBS. Confocal microscopy was performed on a Zeiss confocal laser scanning microscope LSM 700 of the Light Microscopy Facility, a Core Facility of the CMCB Technology Platform at TU Dresden. Antibody used: anti-dIlp7 (gift from Irene Miguel-Aligua, (27))

### GCaMP experiments

#### Sample collection

The egg progeny of crosses between UAS-GCaMP or Canton S and *dilp2-Gal4* or *dilp7-Gal4* were collected. Larvae were prepared as described above in “developmental tracking part”. Three larvae were pipetted in 10μL 0.00005% Triton-X100/PBS solution and transferred in 24 well-plates containing food. Three larvae per well (containing 1mL of food) were deposited. Fed larvae were collected after being reared for 6 or 22 days on yeast food or corn food, respectively.

#### Immunostaining

Performed as described in “Immunochemistry” part. Quantifcation

The total number of dilp2- or dilp7-GCaMP-postive (i.e. green-fluorescent cells) were counted in each brain imaged. The percentage of cells ON was then calculated on the total number of dilp2-positive cells (*i.e.* 8 per hemisphere (14) or dilp7-positive cells (*i.e*. 12 (26)). Statistics, Wilcoxon’s test.

### Larva and adult weight

To determine the weight of larvae or adults reared on yeast food, a determined number of third-instar larvae or adults were put in Eppendorf 2mL tubes. Mass of one larva or one adult (mg/larva or mg/adult, respectively) = (Mass of the tube containing x number of larvae or adults (mT+xL or mT+xA, respectively) – mass of the tube (mT))/x number of larvae or adults contained in the tube (xL or xA, respectively).

### Pupa area

Pupa arising from larvae reared in 96-well plates containing yeast food at 20°C were placed on microscope slides were a 5mm-scale bars were drawn. Pictures were taken with a camera and processed by using Fiji (53). By using the “freehand” tool, the area of each pupa was measured. Kruskal-Wallis tests were performed as statistical tests.

### Feeding behavior

#### Larva mouth contraction

Third-instar larvae fed on normal food were rinsed and transferred to three Petri dishes containing 0.5%Bromophenol-Blue colored yeast food or corn food (27). On each plate, 10 larvae were placed. The number of mouth contraction were counted within 30 seconds for each larva. Kruskal-Wallis tests were performed as statistical tests.

#### Larva food intake

Third-instar larvae fed on normal food were rinsed and transferred to three Petri dishes containing 0.5%Bromophenol-Blue colored yeast food or corn food (27). On each plate, 10 larvae were placed. Every 5 minutes, the number of larvae with blue guts were counted. Finally, the time to reach 100% of the larvae with blue guts per plate was plotted. Kruskal-Wallis tests were performed as statistical tests.

#### Adult food intake

Seven-to-ten days old adults starved for 24 hours were transferred in vials containing 5mL 0.5%Bromophenol-Blue colored yeast food or corn food (27). The male:female ratio per vial was 1:3. Every 30 minutes, the number of females with blue guts were counted. Finally, the percentage of females with blue guts after 4 hours on the diet was plotted. Statistics, experiments out of the range (*i.e.* Mean + or − Standard deviation) were excluded of the normalized data. Kruskal-Wallis tests were performed as statistical tests.

#### Adult transit

Seven-to-ten days old adults kept for 24-48h into vials containing 5mL 0.5%Bromophenol-Blue colored yeast food or corn food (27) were transferred into vials containing starvation diet (25% apple juice, 10%Glucose, 1.5% agar, 0.4% nipagin). The male:female ratio per vial was 1:3. Every 30 minutes, the number of females without blue guts were counted. Finally, the percentage of females without blue guts after 4 hours on starvation was plotted. Statistics, experiments out of the range (*i.e.* Mean + or − Standard deviation) were excluded of the normalized data. Kruskal-Wallis tests were performed as statistical tests.

### Biochemistry

#### AKT detection in yeast and corn-food fed adult heads

Sample preparation, western-blotting and quantification were performed as published by (48). In brief, flies were raised on normal food and 5d old adults transferred onto respective experimental foods for 7d at 20°C. Thereafter, flies were snap-frozen in liquid nitrogen and frozen heads from females collected for processing. Heads were homogenized on ice in 0.01% Tx100-PBS and subsequently cooked at 95°C for 5min. Polyclonal antibodies used to probe were Akt-pSer505 (Cell Signaling, 4054S), Akt-pThr308 (Invitrogen, 44-602G), Akt (Invitrogen, MAS14916), MAPK (Cell Signaling, p44/42 MAPK (Erk1/2) Antibody #9102), P-MAPK (Cell Signaling, Phospho-p44/42 MAPK (Erk1/2) (Thr202/Tyr204) Antibody #9101).

#### Quantification

Pixel intensity of defined area (line) covering the signature (not saturated) center on photograph was measured using FIJI software (53). Intensity ratio between AKT85-Ser505 and panAKT85 or P-MAPK and MAPK was calculated and normalized to value obtained from controls.

### Trehalose, protein and triacylglyceride quantification in larva or adult hemolymph

#### Sample collection and preparation

Larvae: third-instar larvae raised in 96-well plates containing YF were collected, rinsed, drained, transferred in 2ml Eppendorf tubes and short-frozen in liquid nitrogen. Each tube was filled with 20 larvae. To collect hemolymph, 60μL of ice-cold1xPBS was added into the tube containing larvae. After spinning for 5 minutes at 500g, 50μL of the supernatant were pipetted and transferred into another Eppendorf tube.

Adults: eight-to-ten days old adults reared on yeast food were collected, transferred into 2mL Eppendorf tubes and short frozen in liquid nitrogen. For trehalose assay, 460 adults were required (male: female ratio ≈ 230:230); for protein quantification, 16 adults (male: female ratio ≈ 4:12) and for triacylglycerides, 9 adults were needed (male: female ratio ≈ 3:6). To collect hemolymph, 110μL of ice-cold 1xPBS was added to a tube containing sample flies. Samples were incubated on ice for 5 minutes and centrifuged for 10 seconds (benchtop centrifuge) two times, then 100μl of the supernatant was transferred to a fresh tube.…..

#### Trehalose assay and quantification

Hemolymph samples were heated at 85°C for 15 minutes. Then samples were cooled down to room temperature, and trehalose was measured and quantified like recommended by the manufacturer (Trehalose Kit, Megazyme).

#### Protein assay and quantification

Proteins in hemolymph samples were concentrated by choroform-methanol precipitation and re-dissolved in 0.1% TritonX100 in 1xPBS. Protein amount was measured and quantified like recommended by manufacturer (Pierce™ BCA Protein Assay Kit, Thermo Fisher Scientific).

#### Thin layer chromatography (TLCs)

Lipid extraction were performed as described by Bligh and Dyer (54,55) and analyzed on TLC plates using published solvent protocols (polar lipids: CHCl3 : CH3OH : H2O (75 : 25 :2.5); neutral lipids: C7H16:(C2H5)2O :CH3COOH (70 : 30 : 1)) Plates were stained with CuSO4 or Primulin and scanned using a Typhoon reader. Three homogenized fly heads per sample, 1μL lipid extract was pipetted on HPTLC Silica gel glass plates (Merck). Twenty larvae per sample, volume to resupend the lipid pellets was normalized to the larval weight and 3μL lipid extract was pipetted on HPTLC Silica gel glass plates (Merck). Triacylglycerol (TAG) (10mg/ml) and Diacylglycerol (DAG) (1mg/ml) were used as standards.

#### Triacylglycerides quantification

Pixel intensity of defined area (round) covering triglycerides signatures on photograph was measured using FIJI software (53). Intensity was calculated and normalized to the blank value.

## Supporting information

Supporting informations_Raw data

## AKNOWLEDGMENTS

We thank Suzanne Eaton, Irene Miguel-Aliaga and Ronald P. Kühnlein for sharing material. We thank Irene Miguel-Aliaga and her group for their feedback and advices on this project. We thank Mandy Obst for helping with the larva tracking and with lab organization. This work was supported by the Light Microscopy Facility, a Core Facility of the CMCB Technology Platform at TU Dresden. We thank Uenal Coskun’s, Andrej Schevchenko’s and Joerg Mansfeld’s lab members for their help, advice and comments on this project. We thank Julien Marcetteau for his advices and help with figures.

## AUTHOR CONTRIBUTION

Conceptualization: M.B., E.P.; Formal analysis: E.P.; Funding acquisition: M.B.; Investigation: E.P., J.K., L.C.T.; Methodology: E.P.; M.B.; Project administration: M.B., E.P.; Resources: E.P., M.B.; Supervision: E.P., M.B.; Validation: E.P.; Visualization: E.P.; Writing-Original Draft Preparation: M.B., E.P., J.K., L.C.T.

