## Supplementary Figures for "Diet and heat - one neuronal subset two responses"

**SUPPLEMENTARY INFORMATIONS**

Excel sheet showing the raw data for:
- *ΔdIlp* mutant screen on YF, CF at 20°C or 28°C: Pupariation rate (Survival) and developmental speed
- dIlp over-expression on YF, CF at 20°C or 28°C: Pupariation rate (Survival) and developmental speed
- dIlp ectopic expression on YF or CF at 20°C: Pupariation rate (Survival) and developmental speed
- *ΔdIlp* mutant rescue experiments on YF or CF at 20°C: Pupariation rate (Survival) and developmental speed
- *OregonR* larvae on lipid-reduced YF at 20°C or 30°C: Pupariation rate (Survival)
- YF-to-CF larva swap experiment at 20°C: Pupariation rate (Survival)
- IPC or D7N hyperactivation/silencing on YF or CF: Pupariation rate (Survival) and developmental speed
- IPC or D7N silencing induction at different time points on YF: Pupariation rate (Survival) and developmental speed
- GCaMP experiments on YF or CF at 20°C.
- Predicted receptor knock-down on YF at 20°C or 28°C and on CF at 20°C: Pupariation rate (Survival) and developmental speed
- Larva/Adult measurements on YF at 20°C: Larva and adult weight, pupa area
- Larva feeding behavior on YF or CF at 20°C: Mouth hook contractions, food ingestion speed
- Adult feeding behavior on YF or CF at 20°C: Food ingestion and transit
- Larva nutrients on YF at 20°C: Trehalose, protein and TAG level measurements
- Adult nutrients on YF at 20°C or 28°C: Trehalose, protein and TAG level measurements
- Western-blot quantifications

**SUPPLEMENTARY FIGURES**

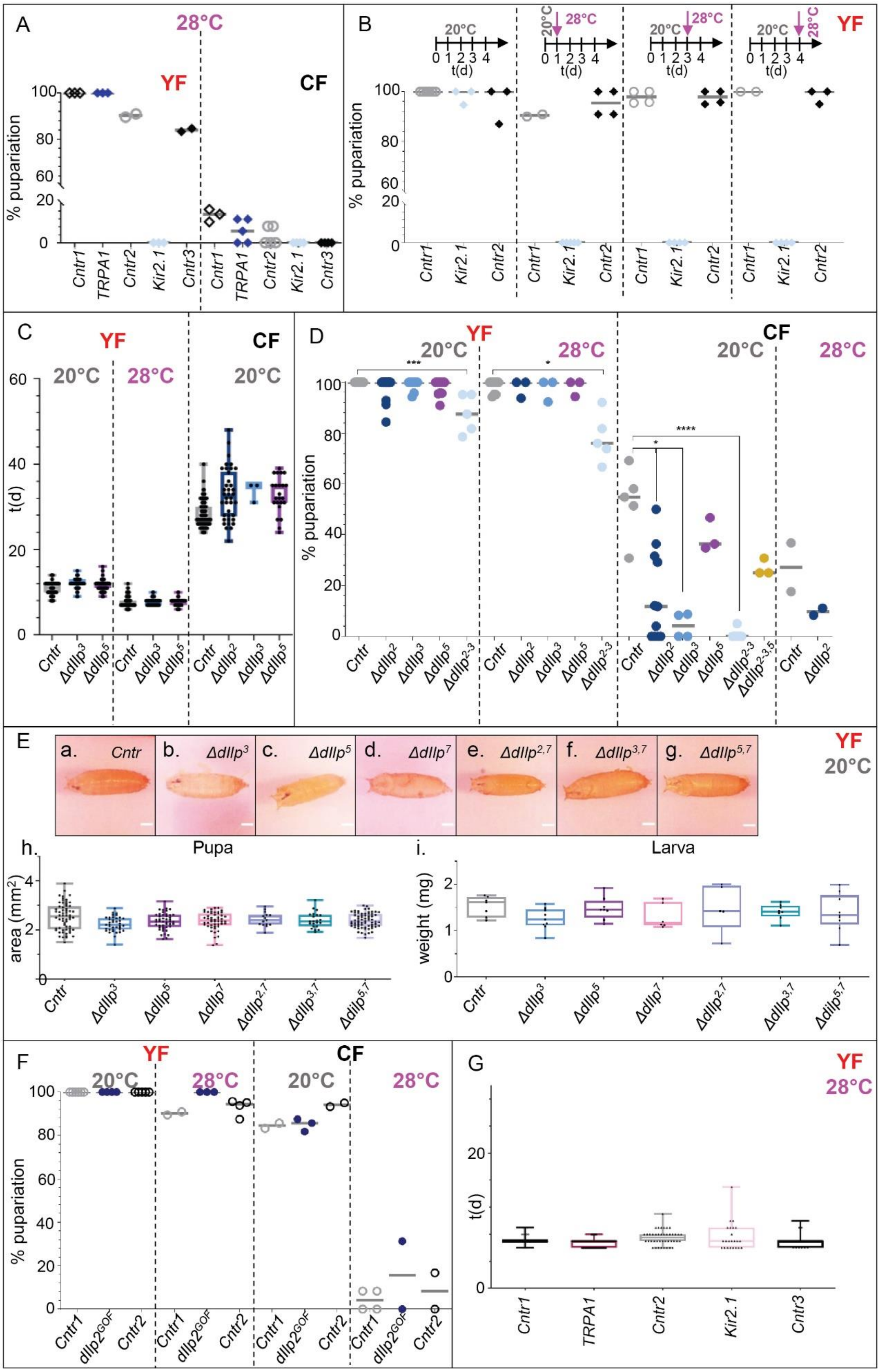

A
28°C
YF
CF
% pupariation
Cntr1
TRPA1
Cntr2
Kir2.1
Cntr3
B
20°C
28°C
t(d)
C
t(d)
Cntr
Δdilp3
Δdilp5
Δdilp2
D
Δdilp2-3
Δdilp2-3,5
E
a. Cntr
b. Δdilp3
c. Δdilp5
d. Δdilp7
e. Δdilp2,7
f. Δdilp3,7
g. Δdilp5,7
h.
Pupa
area (mm2)
i.
Larva
weight (mg)
F
dilp2GOF
G

**Supplementary Figure S1. IPCs and dIlp2, 3 and 5 are required to endure heat stress**
**(A).** Plotted are the pupariation rate (%) of larvae reared on YF (red, left) or CF (black, right) at 28°C. The genotypes shown are the controls *UAS-TRPA1/+* (*Cntr1*, black empty-fill lozenge), *dIlp2-Gal4/+* (*Cntr2*, grey circles), *UAS-Kir2.1,tub80ts/+* (*Cntr3*, black lozenges), *dIlp2>>TRPA1* (*TRPA1*, IPCs hyper-activated, dark-blue lozenges) and *dIlp2>>Kir2.1,tub80ts* (*Kir2.1*, IPCs silenced, light-blue lozenges). One dot=one experiment; the median is depicted as a grey bar. *Cntr1* (YF, median=100%; CF, median=14.3%), *TRPA1* (YF, median=100%; CF, median=5.9%), *Cntr2* (YF, median=90.2%; CF, median= 0%), *Kir2.1* (YF, median=0%; CF, median=0%), *Cntr3* (YF, median=90.2%; CF, median=0%). Each experiment included $n_{larvae}$>6. Statistics, Kruskal-Wallis test p=0.0005 ***, Dunn's multiple comparison test: *Cntr1 vs TRPA1*, $p_{YF}$=0.0575 ns, $p_{CF}$>0.9999 ns; *Cntr2 vs TRPA1*, $p_{YF}$>0.9999 ns, $p_{CF}$=0.2580 ns; *Cntr2 vs Kir2.1*, $p_{YF}$=0.0536 ns, $p_{CF}$=0.3370 ns; *Cntr3 vs Kir2.1*, $p_{YF}$=0.5649 ns, $p_{CF}$>0.9999 ns.
**(B).** Plotted are the pupariation rate (%) of larvae reared on YF and transferred from 20°C to 28°C at different time point (in days, d). From left to right, larvae were raised at 20°C all along development (20°C) or transferred at 28°C after 1 day (1d), 3 days (3d) or 4 days (4d) at 20°C (as explained by scheme on the top). The development of the controls *dilp2-Gal4/+* (*Cntr1*, grey circles), *UAS-Kir2.1, tub80ts/+* (*Cntr2*, dark lozenges) and the *dIlp2>>Kir2.1,tub80ts* (*Kir2.1*, silenced IPCs, light-blue lozenges) was tracked. *UAS-Kir2.1, tub80ts* is an inducible tool *i.e.* at restrictive temperature (20°C), it should not be activated while at permissive temperature (28°) it should be activated in IPCs. Each experiment included $n_{larvae}$ ≥10 (20°C), $n_{larvae}$ ≥6 (1d, 3d and 4d). One dot=one experiment; the median is depicted as a grey bar. *Cntr1* (20°C, median=100%; 1d, median=90.5%; 3d, median=97.9%; 4d, median=100%), *Kir2.1* (20°C, median=100%; 1d, median=0%; 3d, median=0%; 4d, median=0%) and *Cntr2* (20°C, median=100%; 1d, median=95.5%; 3d, median=97.8%; 4d, median=100%). Statistics, Kruskal-Wallis test, p=0.0003 ***; Dunn's multiple comparison test, *Cntr1 vs Kir2.1*: $p_{20°C}$=0.8068 ns, $p_{1d}$>0.3772 ns, $p_{3d}$=0.0331 *, $p_{4d}$=0.0628 ns; *Cntr2 vs Kir2.1*: $p_{20°C}$>0.9999 ns, $p_{1d}$>0.0085 **, $p_{3d}$=0.0581 ns, $p_{4d}$=0.0873 ns.
**(C).** Here is shown the developmental speed (in days, d) of *FOXO*$^{mCherry}$ (*Control*, *Cntr*, grey), *ΔdIlp*$^{3}$ (blue) and *ΔdIlp*$^{5}$ (purple) mutant larvae reared on YF (red) at 20°C or 28°C or on CF (black) at 20°C. *ΔdIlp*$^{2}$ (dark-blue) larvae developmental speed are shown on CF at 20°C (for YF data, see Fig.1A). One black dot = one larva. *Cntr* (YF, median$^{20°C}$=11d; $n_{28}$=169/9, median$^{28°C}$=7d; CF, median$^{20°C}$=27d), *ΔdIlp*$^{3}$ (YF, median$^{20°C}$=12d; median$^{28°C}$=35d), *ΔdIlp*$^{5}$ (YF, median$^{20°C}$ 12d; median$^{28°C}$ =8d; CF, median$^{20°C}$=32d), *ΔdIlp*$^{2}$ (CF, median$^{20°C}$=32.5d). Each experiment included $n_{larvae}$ ≥10 (20°C) or $n_{larvae}$ ≥6 (28°C). Statistics, Kruskal-Wallis test, p<0.0001 ****; Dunn's multiple comparison test, *Cntr vs ΔdIlp*$^{3}$: p$^{20°C}$<0.0001 ****, p$^{28°C}$=0.1055 ns (YF), p$^{20°C}$=0.0606 ns (CF); *Cntr vs ΔdIlp*$^{5}$: p$^{20°C}$=0.0229 *, p$^{28°C}$=0.2331 ns (YF), p$^{20°C}$<0.0001 **** (CF); *Cntr vs ΔdIlp*$^{2}$, p$^{20°C}$<0.0001 **** (CF).
**(D).** Plotted are the pupariation rates (%) of *FOXO*$^{mCherry}$ (*Control*, *Cntr*, grey) and the *ΔdIlp* mutants: *ΔdIlp*$^{2}$ (dark blue), *ΔdIlp*$^{3}$ (blue), *ΔdIlp*$^{5}$ (purple), *ΔdIlp*$^{2\text{-}3}$ (light blue) on YF(red) at 20°C or 28°C or CF at 20°C. Here the data for *ΔdIlp*$^{2\text{-}3,5}$ (yellow) mutant larvae are only shown on CF at 20°C; for the pupariation rates on YF at 20°C or 28°C, see the Fig. 2B or Fig. 1C, respectively. On CF, at 28°C, only the development of *ΔdIlp*$^{2}$ mutants was analysed. *Cntr* (YF, median$^{20°C}$=100%, median$^{28°C}$=100%; CF, median$^{20°C}$=54.8%, median$^{28°C}$=27.2%), *ΔdIlp*$^{2}$ (YF, median$^{20°C}$=100%, median$^{28°C}$=100%; CF, median$^{20°C}$=11.8%, median$^{28°C}$=9.7%), *ΔdIlp*$^{3}$ (YF, median$^{20°C}$=100%, median$^{28°C}$=100%; CF, median$^{20°C}$=4.2%), *ΔdIlp*$^{5}$ (YF, median$^{20°C}$=100%, median$^{28°C}$=100%; CF, median$^{20°C}$=36.4%), *ΔdIlp*$^{2\text{-}3}$ (YF, median$^{20°C}$=84.7%, median$^{28°C}$=78.9%; CF, median$^{20°C}$=0%), *ΔdIlp*$^{2\text{-}3,5}$ (CF, median$^{20°C}$=27.9%). Statistics, Kruskal-Wallis test, p<0.0001 ****. Dunn's multiple comparison test, *Cntr vs ΔdIlp*$^{2}$: p$^{20°C}$>0.9999 ns, p$^{28°C}$>0.9999 ns (YF), p$^{20°C}$=0.0430 * (CF); *Cntr vs ΔdIlp*$^{3}$: p$^{20°C}$>0.9999 ns, p$^{28°C}$>0.9999 ns (YF), p$^{20°C}$=0.0214 * (CF); *Cntr vs ΔdIlp*$^{5}$: p$^{20°C}$=0.5747 ns, p$^{28°C}$>0.9999 ns (YF), p$^{20°C}$>0.9999 (CF); *Cntr vs ΔdIlp*$^{2\text{-}3}$: p$^{20°C}$=0.0004 ***, p$^{28°C}$=0.0028 * (YF), p$^{20°C}$<0.0001 **** (CF). *Cntr vs ΔdIlp*$^{2\text{-}3,5}$: p$^{20°C}$>0.9999 ns (CF). Mann-Whitney test, *Cntr vs ΔdIlp*$^{2}$: p$^{28°C}$=0.3333 ns (CF).
**(E). a-h.** Representative pictures of pupae reared on YF (a-g) whose area has been measured (Fig. 1Ba) and plotted (h). The area (in $mm^2$) has been analysed for *FOXO*$^{mCherry}$ (a, h; *Control*, *Cntr*), *ΔdIlp*$^{3}$ (b), *ΔdIlp*$^{5}$ (c), *ΔdIlp*$^{7}$ (d), *ΔdIlp*$^{2,7}$ (e), *ΔdIlp*$^{3,7}$ (f) and *ΔdIlp*$^{5,7}$(g). **h.** One black dot=one pupa. *Cntr* (grey, median=2.56), *ΔdIlp*$^{3}$ (blue, median=2.22), *ΔdIlp*$^{5}$ (purple, median=2.34), *ΔdIlp*$^{7}$ (pink, median=2.39), *ΔdIlp*$^{2,7}$ (blue, median=2.40), *ΔdIlp*$^{3,7}$ (green-blue, median=2.36) and *ΔdIlp*$^{5,7}$ (light-purple, median=2.36). Statistics, Kruskal-Wallis test, p=0.0770 ns. Dunn's multiple comparison test, *Cntr vs*

*ΔdIlp*$^{3}$: p=0.0062 **, *Cntr vs ΔdIlp*$^{5}$: p=0.3405 ns, *Cntr vs ΔdIlp*$^{7}$, *ΔdIlp*$^{2,7}$ or *ΔdIlp*$^{3,7}$: p>0.9999 ns; *Cntr vs ΔdIlp*$^{5,7}$: p=0.7383 ns. **i.** The weight (in mg per larva) was calculated from n larva pools (n-p). Each pool contained $n_{larvae}$ ≥8. *Cntr* (grey, median=1.61), *ΔdIlp*$^{3}$ (blue, median=1.24), *ΔdIlp*$^{5}$ (purple, median=1.45), *ΔdIlp*$^{7}$ (pink, median=1.16), *ΔdIlp*$^{2,7}$ (blue, median=1.42), *ΔdIlp*$^{3,7}$ (green-blue, median=1.41) and *ΔdIlp*$^{5,7}$ (light purple, median=1.33). Statistics, Kruskal-Wallis test, p=0.3856 ns. Dunn's multiple comparison test, *Cntr vs ΔdIlp*$^{3}$: p=0.3405 ns, *Cntr vs ΔdIlp*$^{5}$, *ΔdIlp*$^{2,7}$, *ΔdIlp*$^{3,7}$ or *ΔdIlp*$^{5,7}$: p>0.9999 ns, *Cntr vs ΔdIlp*$^{7}$: p=0.5978 ns.

**(F).** Here are plotted the pupariation rates (%) of the controls *dIlp2-Gal4/+* (*Control1*, *Cntr1*, grey circles), *UAS-DIlp2/+* (*Control2*, *Cntr2*, black circles) and the dIlp2 gain-of-function (*dIlp2*$^{GOF}$, *dIlp2>>dIlp2*, blue rounds) larvae kept on YF (red) or CF (black) at 20°C or 28°C. One dot=one experiment; the median is depicted as a grey bar. Each experiment included $n_{larvae}$>10 (20°C) or $n_{larvae}$>6 (28°C). *Cntr1* (YF, $\text{median}^{20°C}$=100%, $\text{median}^{28°C}$=90.2%; CF, $\text{median}^{20°C}$=84.5%, $\text{median}^{28°C}$=4.2%); *dIlp2*$^{GOF}$ (YF, $\text{median}^{20°C}$=100%, $\text{median}^{28°C}$=100%; CF, $\text{median}^{20°C}$=85.7%, $\text{median}^{28°C}$=15.6%) and *Cntr2* (YF, $\text{median}^{20°C}$=100%, $\text{median}^{28°C}$=94.5%; CF, $\text{median}^{20°C}$=94.2%, $\text{median}^{28°C}$=8.3%). Statistics, Kruskal-Wallis test, p<0.0001 ****. Dunn's multiple comparison test, *Cntr1 vs dIlp2*$^{GOF}$: $p^{20°C}$>0.9999 ns, $p^{28°C}$=0.0757 ns (YF), $p^{20°C}$>0.9999 ns, $p^{28°C}$>0.9999 ns (CF); *Cntr2 vs dIlp2*$^{GOF}$: $p^{20°C}$>0.9999 ns, $p^{28°C}$>0.9999 ns (YF), $p^{20°C}$=0.2642 ns, $p^{28°C}$>0.9999 ns (CF).

**(G).** Here is shown the developmental speed (in days, d) of *UAS-TRPA1/+* (*Control*, *Cntr1*, black, black empty-fill lozenges), *dIlp7>>TRPA1* (*TRPA1*, hyper-activated D7Ns, dark pink), *dIlp7-Gal4/+* (*Control*, *Cntr2*, grey), *dIlp7>>Kir2.1,tub80ts* (*Kir2.1*, silenced D7Ns, dark pink) and *UAS-Kir2.1,tub80ts/+* (*Control*, *Cntr3*, black, black lozenges) larvae reared on YF at 28°C. One black dot = one larva. *Cntr1* (median=7d), *TRPA1* (median=7d), *Cntr2* (median=7.5d), *Kir2.1* (median=7d) and *Cntr3* (median=7d). Each experiment included $n_{larvae}$ ≥6. Statistics, Kruskal-Wallis test, p=0.0002 ***. Dunn's multiple comparison test, *Cntr1 vs TRPA1*: p<0.0001 ****; *Cntr2 vs TRPA1*: p=0.0051 ** ; *Cntr2 vs Kir2.1*: p>0.9999 ns; *Cntr3 vs Kir2.1*: p>0.9999 ns.

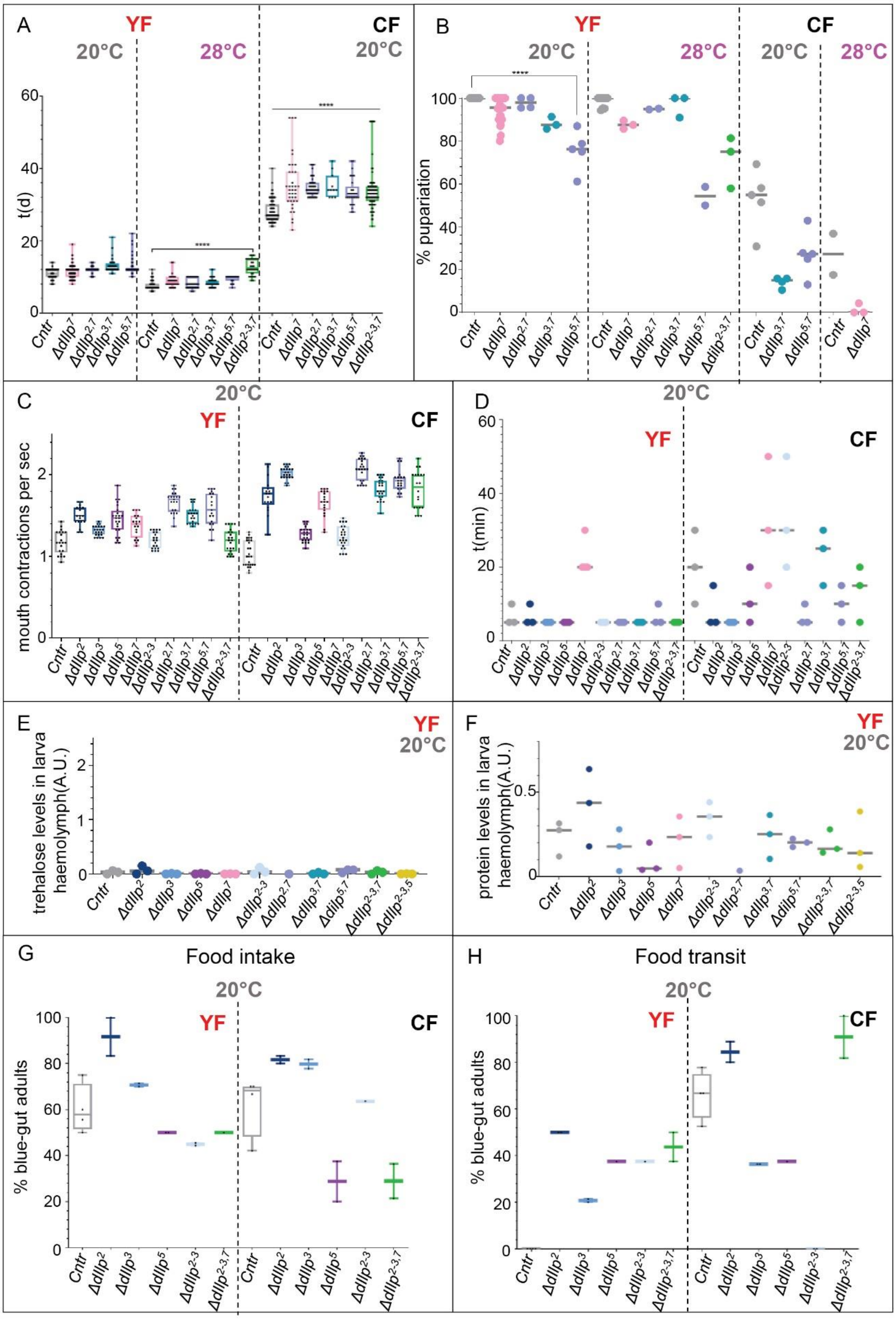

A
YF
20°C
28°C
CF
20°C
t(d)
60
40
20
0
****
****
Cntr
ΔdIlp7
ΔdIlp2,7
ΔdIlp3,7
ΔdIlp5,7
ΔdIlp2-3,7
B
YF
20°C
28°C
CF
20°C
28°C
% pupariation
100
80
60
40
20
0
****
C
20°C
YF
CF
mouth contractions per sec
2
1
0
ΔdIlp2
ΔdIlp3
ΔdIlp5
ΔdIlp2-3
D
20°C
YF
CF
t(min)
60
40
20
0
E
YF
20°C
trehalose levels in larva haemolymph(A.U.)
2
1
0
F
YF
20°C
protein levels in larva haemolymph(A.U.)
0.5
0
ΔdIlp2-3,5
G
Food intake
20°C
YF
CF
% blue-gut adults
100
80
60
40
20
0
H
Food transit
20°C
YF
CF
% blue-gut adults
100
80
60
40
20
0

**Supplementary Figure S2. DIlp7 peptide is instructive for larval feeding behavior on yeast-free diet**

**(A).** Plotted is the developmental speed (in days, d) of larvae reared on YF (red, left) at 20°C or 28°C, or on CF (black, right) at 20°C. *FOXO*$^{mCherry}$ (*Control*, *Cntr*, grey), *ΔdIlp*$^{7}$ (pink), *ΔdIlp*$^{2,7}$ (blue), *ΔdIlp*$^{3,7}$ (green-blue), *ΔdIlp*$^{5,7}$ (light purple) and *ΔdIlp*$^{2\text{-}3,7}$ (green) larvae were tracked. For the plot of the developmental speed of *ΔdIlp*$^{2\text{-}3,7}$ larvae on YF at 20°C, see Fig. 2D. One black dot = one larva. *Cntr* (YF, median$^{20°C}$=11d, median$^{28°C}$=7d; CF, median$^{20°C}$=27d), *ΔdIlp*$^{7}$ (YF, median$^{20°C}$=12d, median$^{28°C}$=8.5d; CF, median$^{20°C}$=34d), *ΔdIlp*$^{2,7}$ (YF, median$^{20°C}$=12d, median$^{28°C}$=8d; CF, median$^{20°C}$=34d), *ΔdIlp*$^{3,7}$ (YF, median$^{20°C}$=13d, median$^{28°C}$=8d; CF, n median$^{20°C}$=34d), *ΔdIlp*$^{5,7}$ (YF, median$^{20°C}$=12d, median$^{28°C}$ =1d; CF, median$^{20°C}$=33d) and *ΔdIlp*$^{2\text{-}3,7}$ larvae (YF, median$^{28°C}$=12.5d; CF, median$^{20°C}$=33d). Each experiment included $n_{larvae} \geq 10$ (20°C) or $n_{larvae} \geq 6$ (28°C). Statistics, Kruskal-Wallis test, p<0.0001 ****. Dunn's multiple comparison test, *Cntr* vs *ΔdIlp*$^{7}$: p$^{20°C}$= 0.0517 ns, p$^{28°C}$= 0.0002 *** (YF), p$^{20°C}$<0.0001 **** (CF); *Cntr* vs *ΔdIlp*$^{2,7}$: p$^{20°C}$<0.0001 ****, p$^{28°C}$= 0.1742 ns (YF), p$^{20°C}$<0.0001 **** (CF); *Cntr* vs *ΔdIlp*$^{3,7}$: p$^{20°C}$<0.0001 ****, p$^{28°C}$= 0.0005 *** (YF), p$^{20°C}$<0.0001 **** (CF); *Cntr* vs *ΔdIlp*$^{5,7}$: p$^{20°C}$<0.0001 ****, p$^{28°C}$<0.0001 **** (YF), p$^{20°C}$<0.0001 **** (CF); *Cntr* vs *ΔdIlp*$^{2\text{-}3,7}$: p$^{28°C}$<0.0001 **** (YF), p$^{20°C}$<0.0001 **** (CF).

**(B).** Plotted are the pupariation rates (%) of larvae reared on YF (red, left) or on CF (black, right) at 20°C or 28°C. *FOXO*$^{mCherry}$ (*Control*, *Cntr*, grey), *ΔdIlp*$^{7}$ (pink), *ΔdIlp*$^{2,7}$ (blue), *ΔdIlp*$^{3,7}$ (green-blue), *ΔdIlp*$^{5,7}$ (light purple) and *ΔdIlp*$^{2\text{-}3,7}$ (green) larvae were tracked. On CF, at 28°C, only the development of *ΔdIlp*$^{7}$ has been tracked and analysed. For the plot depicting the pupariation rate of *ΔdIlp*$^{7}$ larvae on CF at 20°C, see Fig. 2A; for *ΔdIlp*$^{2\text{-}3,7}$ larvae kept on YF or CF at 20°C, see Fig. 2B or Fig. 2A, respectively. One dot =one larva; the median is shown as a grey bar. Each experiment included $n_{larvae}>10$ (20°C) or $n_{larvae}>6$ (28°C). *Cntr* (YF, median$^{20°C}$=100%, median$^{28°C}$=100%; CF, median$^{20°C}$=54.8%, median$^{28°C}$=27.2%), *ΔdIlp*$^{7}$ (YF, median$^{20°C}$=95.5%, median$^{28°C}$=87.5%; CF, median$^{28°C}$=0%), *ΔdIlp*$^{2,7}$ (YF, median$^{20°C}$=97.8%, median$^{28°C}$=95.0%), *ΔdIlp*$^{3,7}$ (YF, median$^{20°C}$=87.5%, median$^{28°C}$=95.5%; CF, median$^{20°C}$=15.0%), *ΔdIlp*$^{5,7}$ (YF, median$^{20°C}$=76.2%, median$^{28°C}$=54.3%; CF, median$^{20°C}$=27.3%), *ΔdIlp*$^{2\text{-}3,7}$ (YF, median$^{28°C}$=78.1%). Statistics, Kruskal-Wallis test, p<0.0001 ****. Dunn's multiple comparison test, *Cntr* vs *ΔdIlp*$^{7}$: p$^{20°C}$=0.0971 ns, p$^{28°C}$=0.1311 ns (YF); *Cntr* vs *ΔdIlp*$^{2,7}$: p$^{20°C}$>0.9999 ns, p$^{28°C}$>0.9999 ns (YF); *Cntr* vs *ΔdIlp*$^{3,7}$: p$^{20°C}$=0.0178 *, p$^{28°C}$>0.9999 ns (YF), p$^{20°C}$=0.0046 ** (CF); *Cntr* vs *ΔdIlp*$^{5,7}$: p$^{20°C}$<0.0001 ****, p$^{28°C}$=0.0184 * (YF), p$^{20°C}$=0.0982 ns (CF); *Cntr* vs *ΔdIlp*$^{2\text{-}3,7}$: p$^{28°C}$=0.0130 * (YF). Mann-Whitney test, *Cntr* vs *ΔdIlp*$^{7}$: p$^{28°C}$=0.2000 ns (CF).

**(C).** Plotted are the number of mouth contractions per seconds of larvae put on YF (red, left) or on CF (black, right) at 20°C. *FOXO*$^{mCherry}$ (*Control*, *Cntr*, grey) and *ΔdIlp* mutant larvae were analysed. One black dot = one larva. For each genotype, $N_{exp}$= 3. *Cntr* (YF, median=1.17; CF, median=1.00), *ΔdIlp*$^{2}$ (dark blue; YF, median=1.50; CF, median=1.77), *ΔdIlp*$^{3}$ (blue; YF, median=1.33; CF, median= 2.03), *ΔdIlp*$^{5}$ (dark purple; YF, median=1.47; CF, median= 1.28), *ΔdIlp*$^{7}$ (pink; YF, median=1.40; CF, median=1.67), *ΔdIlp*$^{2\text{-}3}$ (light blue; YF, median=1.20; CF, median=1.23), *ΔdIlp*$^{2,7}$ (blue; YF, median=1.70; CF, median=2.07), *ΔdIlp*$^{3,7}$ (green-blue; YF, median=1.57; CF, median=1.80), *ΔdIlp*$^{5,7}$ (light purple; YF, median=1.57; CF, median=1.93), *ΔdIlp*$^{2\text{-}3,7}$ (green; YF, median=1.20; CF, median=1.85). Statistics, Kruskal-Wallis test, p<0.0001 ****. Dunn's multiple comparison test, *Cntr* vs *ΔdIlp*$^{2}$: $p_{YF}$, $p_{CF}$<0.0001 ****; *Cntr* vs *ΔdIlp*$^{3}$: $p_{YF}$=0.3564 ns, $p_{CF}$<0.0001 ****; *Cntr* vs *ΔdIlp*$^{5}$: $p_{YF}$<0.0001 ****, $p_{CF}$=0.9662 ns; *Cntr* vs *ΔdIlp*$^{7}$: $p_{YF}$=0.0147 *, $p_{CF}$=0.0003 ***; *Cntr* vs *ΔdIlp*$^{2\text{-}3}$: $p_{YF}$, $p_{CF}$>0.9999 ns; *Cntr* vs *ΔdIlp*$^{2,7}$: $p_{YF}$, $p_{CF}$<0.0001 ****; *Cntr* vs *ΔdIlp*$^{3,7}$: $p_{YF}$, $p_{CF}$<0.0001 ****; *Cntr* vs *ΔdIlp*$^{5,7}$: $p_{YF}$, $p_{CF}$<0.0001 ****; *Cntr* vs *ΔdIlp*$^{2\text{-}3,7}$: $p_{YF}$>0.9999 ns, $p_{CF}$<0.0001 ****.

**(D).** Here is shown the time (in min) to get 100% of the larvae with blue gut after their transfer on blue-stained YF (red, left) or CF (black, right) at 20°C. *FOXO*$^{mCherry}$ (*Control*, *Cntr*, grey) and *ΔdIlp* mutant larvae were analysed. One black dot = one replicate. For each genotype, three replicates with 10 larvae were performed. *Cntr* ($median_{YF}$=5, $median_{CF}$=20), *ΔdIlp*$^{2}$ (dark blue, $median_{YF}$=5, $median_{CF}$=5), *ΔdIlp*$^{3}$ (blue, $median_{YF}$=5, $median_{CF}$=5), *ΔdIlp*$^{5}$ (dark purple, $median_{YF}$=5, $median_{CF}$=10), *ΔdIlp*$^{7}$ (pink, $median_{YF}$=20, $median_{CF}$=30), *ΔdIlp*$^{2\text{-}3}$ (light blue, $median_{YF}$=5, $median_{CF}$=30), *ΔdIlp*$^{2,7}$ (blue, $median_{YF}$=5, $median_{CF}$=5), *ΔdIlp*$^{3,7}$ (green-blue, $median_{YF}$=5, $median_{CF}$=25), *ΔdIlp*$^{5,7}$ (light purple, $median_{YF}$=5, $median_{CF}$=10), *ΔdIlp*$^{2\text{-}3,7}$ (green, $median_{YF}$=5, $median_{CF}$=15). Statistics, Kruskal-Wallis test, p=0.0005 ***. Dunnett's multiple comparison test, *Cntr* vs *ΔdIlp*$^{2}$: $p_{YF}$, $p_{CF}$>0.9999 ns; *Cntr* vs *ΔdIlp*$^{3}$: $p_{YF}$<0.9999 ns $p_{CF}$=0.3285 ns; *Cntr* vs *ΔdIlp*$^{5}$: $p_{YF}$, $p_{CF}$>0.9999 ns; *Cntr* vs *ΔdIlp*$^{7}$: $p_{YF}$=0.1214 ns, $p_{CF}$>0.9999 ns; *Cntr* vs *ΔdIlp*$^{2\text{-}3}$: $p_{YF}$, $p_{CF}$>0.9999 ns; *Cntr* vs *ΔdIlp*$^{2,7}$: $p_{YF}$<0.9999 ns, $p_{CF}$=0.7078 ns; *Cntr* vs *ΔdIlp*$^{3,7}$: $p_{YF}$, $p_{CF}>0.9999$ ns; *Cntr* vs *ΔdIlp*$^{5,7}$: $p_{YF}$, $p_{CF}>0.9999$ ns; *Cntr* vs *ΔdIlp*$^{2\text{-}3,7}$: $p_{YF}$, $p_{CF}>0.9999$ ns.

**(E).** Here is shown the trehalose levels (in arbitrary units, A.U.) in larva haemolymph. *FOXO*$^{mCherry}$ (*Control, Cntr,* grey), *ΔdIlp*$^{2}$ (dark blue), *ΔdIlp*$^{3}$ (blue), *ΔdIlp*$^{5}$ (purple), *ΔdIlp*$^{7}$ (purple), *ΔdIlp*$^{2\text{-}3}$ (light blue), *ΔdIlp*$^{2,7}$ (blue), *ΔdIlp*$^{3,7}$ (blue)and *ΔdIlp*$^{2\text{-}3,7}$ (green) larvae were grown on YF at 20°C. *Cntr* (median=0.033), *ΔdIlp*$^{2}$ (median=0.055), *ΔdIlp*$^{3}$ (median=0.000), *ΔdIlp*$^{5}$ (median=0.000), *ΔdIlp*$^{7}$ (median=0.000), *ΔdIlp*$^{2\text{-}3}$ (median=0.035), *ΔdIlp*$^{2,7}$ (median=0.000), *ΔdIlp*$^{3,7}$ (median=0.000), *ΔdIlp*$^{5,7}$ (median=0.076), *ΔdIlp*$^{2\text{-}3,7}$ (median=0.030) and *ΔdIlp*$^{2\text{-}3,5}$ (median=0.000).Statistics, Kruskal-Wallis, p=0.2924 ns. Dunn's multiple comparison test, *Cntr* vs *ΔdIlps*: p>0.9999 ns.

**(F).** Here is shown the protein levels (in arbitrary units, A.U.) in larva haemolymph. *FOXO*$^{mCherry}$ (*Control, Cntr,* grey), *ΔdIlp*$^{2}$ (dark blue), *ΔdIlp*$^{3}$ (blue), *ΔdIlp*$^{5}$ (purple), *ΔdIlp*$^{7}$ (purple), *ΔdIlp*$^{2\text{-}3}$ (light blue), *ΔdIlp*$^{2,7}$ (blue), *ΔdIlp*$^{3,7}$ (blue)and *ΔdIlp*$^{2\text{-}3,7}$ (green) larvae were grown on YF at 20°C. *Cntr* (median=0.275), *ΔdIlp*$^{2}$ (median=0.437), *ΔdIlp*$^{3}$ (median=0.179), *ΔdIlp*$^{5}$ (median=0.048), *ΔdIlp*$^{7}$ (median=0.234), *ΔdIlp*$^{2\text{-}3}$ (median=0.356), *ΔdIlp*$^{2,7}$ (median=0.035), *ΔdIlp*$^{3,7}$ (median=0.252), *ΔdIlp*$^{5,7}$ (median=0.202), *ΔdIlp*$^{2\text{-}3,7}$ (median=0.165) and *ΔdIlp*$^{2\text{-}3,5}$ (median=0.140). Statistics, Kruskal-Wallis, p=0.2924 ns. Dunn's multiple comparison test, *Cntr* vs *ΔdIlps*: p>0.9999 ns.

**(G).** Here is plotted the percentage of *CantonS* (Control, *Cntr*, grey), *ΔdIlp*$^{2}$ (dark blue), *ΔdIlp*$^{3}$ (blue), *ΔdIlp*$^{5}$ (purple), *ΔdIlp*$^{2\text{-}3}$ (light blue) and *ΔdIlp*$^{2\text{-}3,7}$ (green) adult females with blue gut, 4 hours after their transfer from uncoloured starvation plates to blue-stained YF (red, left) or CF (black, right). One black dot = one experiment. *Cntr* (YF, median=57.80%; CF, median=68.35%), *ΔdIlp*$^{2}$ (YF, median=91.7%; CF, median=81.7%), *ΔdIlp*$^{3}$ (YF, median=70.7%; CF, median=79.8%), *ΔdIlp*$^{5}$ (YF, median=50.0%; CF, median=28.8%), *ΔdIlp*$^{2\text{-}3}$ (YF, median=44.9%; CF, median=63.6%) and *ΔdIlp*$^{2\text{-}3,7}$ (YF, median=50.0%; CF, median=28.9%). Each experiment included $n_{females} \geq 6$. Statistics, Kruskal-Wallis test, p=0.0242 *. Dunn's multiple comparison test, *Cntr vs ΔdIlp*$^{2}$ : $p_{YF}=0.7136$ ns, $p_{CF}=0.7922$ ns; *Cntr vs ΔdIlp*$^{3}$ : $p_{YF}$, $p_{CF}>0.9999$ ns; *Cntr vs ΔdIlp*$^{5}$ : $p_{YF}>0.9999$ ns, $p_{CF}=0.7922$ ns; *Cntr vs ΔdIlp*$^{2\text{-}3}$ : $p_{YF}=0.3277$ ns, $p_{CF}>0.9999$ ns; *Cntr vs ΔdIlp*$^{2\text{-}3,7}$ : $p_{YF}>0.9999$ ns, $p_{CF}=0.7922$ ns.

**(H).** Here is plotted the percentage of *CantonS* (Control, *Cntr*, grey), *ΔdIlp*$^{2}$ (dark blue), *ΔdIlp*$^{3}$ (blue), *ΔdIlp*$^{5}$ (purple), *ΔdIlp*$^{2\text{-}3}$ (light blue) and *ΔdIlp*$^{2\text{-}3,7}$ (green) adult females with blue gut 4 hours after their transfer from blue-stained YF (red, left) or CF (black, right) to uncoloured starvation plates. One black dot = one experiment. *Cntr* (YF, median=0%; CF, median=66.70%), *ΔdIlp*$^{2}$ (YF, median=50.0%; CF, median=84.4%), *ΔdIlp*$^{3}$ (YF, median=20.7%; CF, median=36.4%), *ΔdIlp*$^{5}$ (YF, median=37.5%; CF, median=37.5%), *ΔdIlp*$^{2\text{-}3}$ (YF, median=37.5%; CF, median=0%) and *ΔdIlp*$^{2\text{-}3,7}$ (YF, median=43.8%; CF, median=90.9%). Each experiment included $n_{females} \geq 3$. Statistics, Kruskal-Wallis test, p=0.0140 *. Dunn's multiple comparison test, *Cntr vs ΔdIlp*$^{2}$: $p_{YF}=0.0246$ *, $p_{CF}>0.9999$ ns; *Cntr vs ΔdIlp*$^{3}$ : $p_{YF}$, $p_{CF}>0.9999$ ns; *Cntr vs ΔdIlp*$^{5}$ : $p_{YF}=0.7935$ ns, $p_{CF}>0.9999$ ns; *Cntr vs ΔdIlp*$^{2\text{-}3}$ : $p_{YF}=0.7935$ ns, $p_{CF}>0.9999$ ns; *Cntr vs ΔdIlp*$^{2\text{-}3,7}$ : $p_{YF}=0.1028$ ns, $p_{CF}=0.9016$ ns.

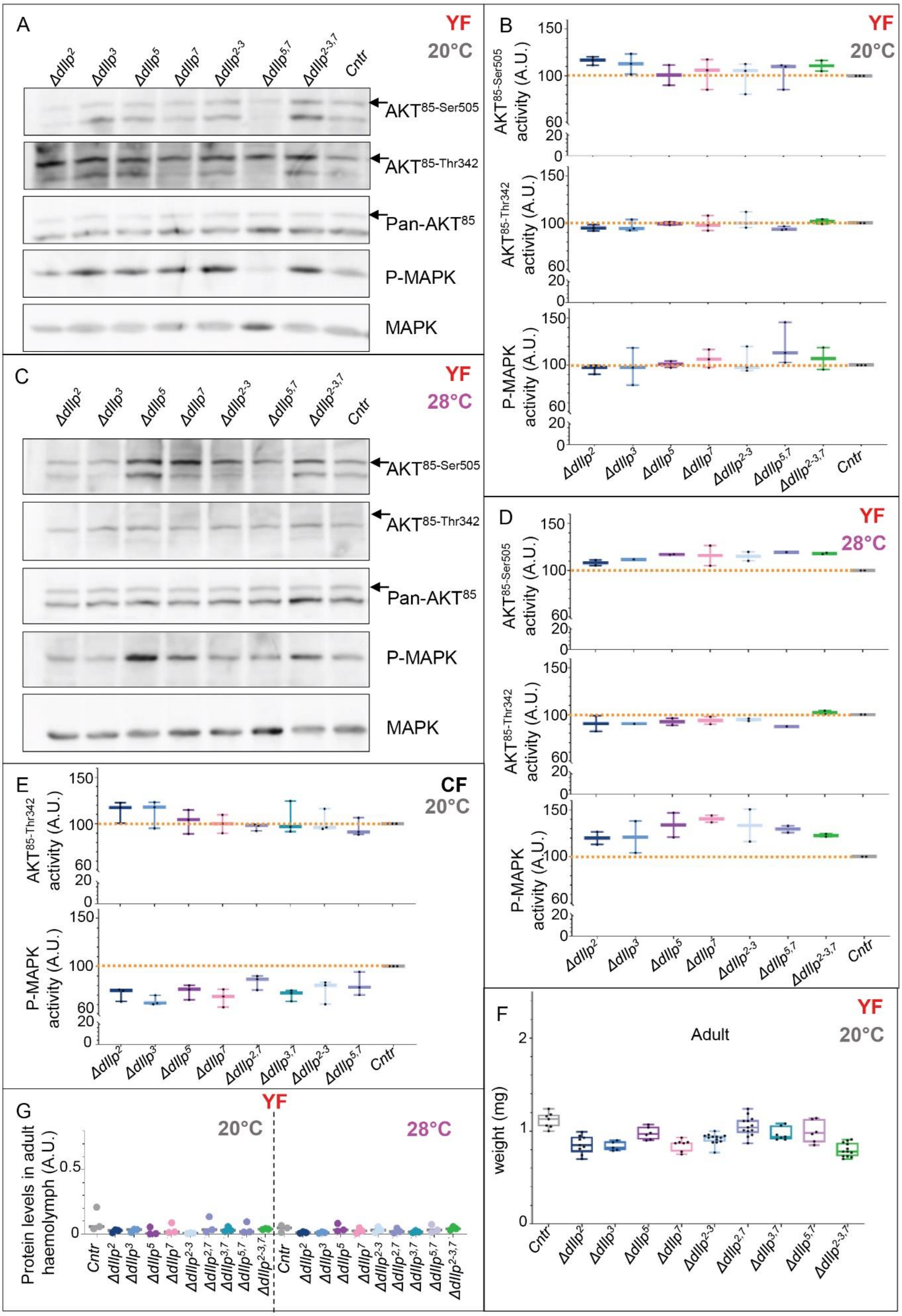
A
YF
20°C
Δdilp2 Δdilp3 Δdilp5 Δdilp7 Δdilp2-3 Δdilp5,7 Δdilp2-3,7 Cntr
AKT85-Ser505
AKT85-Thr342
Pan-AKT85
P-MAPK
MAPK
B
YF
20°C
AKT85-Ser505 activity (A.U.)
AKT85-Thr342 activity (A.U.)
P-MAPK activity (A.U.)
C
YF
28°C
D
YF
28°C
E
CF
20°C
Δdilp2 Δdilp3 Δdilp5 Δdilp7 Δdilp2,7 Δdilp3,7 Δdilp2-3 Δdilp5,7 Cntr
F
YF
Adult
20°C
weight (mg)
Cntr Δdilp2 Δdilp3 Δdilp5 Δdilp7 Δdilp2-3 Δdilp2,7 Δdilp3,7 Δdilp5,7 Δdilp2-3,7
G
YF
20°C
28°C
Protein levels in adult haemolymph (A.U.)

**Supplementary Figure S3. Activity of anabolic and mitogenic IS in adults**

**(A-E).** The phosphorylation state of AKT and MAPK has been analysed by western-blotting (A, C) and quantified (B, D, E) for adults fed on YF at 20°C (A, B) or 28°C (C, D), or on CF at 20°C (E). **A, C.** In adults, two AKT isoforms can be detected, $AKT^{66}$ (66KDa) and $AKT^{85}$ (85KDa); only $AKT^{85}$ has been analysed (shown by a black arrow). AKT can be phosphorylated on two positions, $Serin^{505}$ ($AKT^{85\text{-}Ser505}$) and $Threonin^{342}$ ($AKT^{85\text{-}Thr342}$). The phosphorylated form of MAPK is detected by P-MAPK. The total amount of $AKT^{85}$ or MAPK are shown with Pan-AKT or MAPK, respectively. The blots for adults fed on CF at 20°C are shown in Fig.3C. The genotypes are written above the blots. **B, D, E.** The activity of $AKT^{85\text{-}Ser505}$, $AKT^{85\text{-}Thr342}$ or P-MAPK (in arbitrary units, A.U.) has been quantified and plotted for adults fed on YF at 20°C (B), 28°C (D) or on CF at 20°C (E). The quantifications were performed on n western-blot (n-w). One black dot = 1 n-w. For the quantification of $AKT^{85\text{-}Ser505}$ in animals fed on CF, see Fig. 3C. The activity of each genotype was normalized to the control $FOXO^{mCherry}$ (*Cntr*, grey, median=100%). **B.** *Graphs from top to bottom.* $AKT^{85\text{-}Ser505}$: $\Delta dIlp^{2}$ (dark blue, n-w=3, median=116.7), $\Delta dIlp^{3}$ (blue, n-w=3, median=113.0), $\Delta dIlp^{5}$ (dark purple, n-w=2, median=100.9), $\Delta dIlp^{7}$ (pink, n-w=3, median=106.1), $\Delta dIlp^{2\text{-}3}$ (light blue, n-w=3, median=105.5), $\Delta dIlp^{5,7}$ (light purple, n-w=3, median=109.8), $\Delta dIlp^{2\text{-}3,7}$ (green, n-w=2, median=110.8), *Cntr* (n-w=3). Statistics, Kruskal-Wallis test, p=0.3853 ns. Dunn's multiple comparison test, *Cntr* vs $\Delta dIlp^{2}$: p=0.1776 ns; *Cntr* vs $\Delta dIlp^{3}$: p=0.4746 ns; *Cntr* vs $\Delta dIlp^{5}$, $\Delta dIlp^{7}$, $\Delta dIlp^{2\text{-}3}$, $\Delta dIlp^{5,7}$, $\Delta dIlp^{2\text{-}3,7}$: p>0.9999 ns. $AKT^{85\text{-}Thr342}$: $\Delta dIlp^{2}$ (dark blue, n-w=3, median=94.6), $\Delta dIlp^{3}$ (blue, n-w=3, median=94.1), $\Delta dIlp^{5}$ (dark purple, n-w=2, median=99.5), $\Delta dIlp^{7}$ (pink, n-w=3, median=97.6), $\Delta dIlp^{2\text{-}3}$ (light blue, n-w=3, median=99.6), $\Delta dIlp^{5,7}$ (light purple, n-w=3, median=93.4), $\Delta dIlp^{2\text{-}3,7}$ (green, n-w=2, median=101.6), *Cntr* (n-w=3). Statistics, Kruskal-Wallis test, p=0.3370 ns. Dunn's multiple comparison test, *Cntr* vs $\Delta dIlp^{2}$: p=0.5461 ns; Cntr vs $\Delta dIlp^{5,7}$: p=0.4130 ns; *Cntr* vs $\Delta dIlp^{3}$, $\Delta dIlp^{5}$, $\Delta dIlp^{7}$, $\Delta dIlp^{2\text{-}3}$, $\Delta dIlp^{2\text{-}3,7}$: p>0.9999 ns. P-MAPK: $\Delta dIlp^{2}$ (dark blue, n-w=3, median=97.4), $\Delta dIlp^{3}$ (blue, n-w=3, median=97.4), $\Delta dIlp^{5}$ (dark purple, n-w=2, median=100.9), $\Delta dIlp^{7}$ (pink, n-w=3, median=106.4), $\Delta dIlp^{2\text{-}3}$ (light blue, n-w=3, median=97.6), $\Delta dIlp^{5,7}$ (light purple, n-w=3, median=113.2), $\Delta dIlp^{2\text{-}3,7}$ (green, n-w=2, median=107.1), *Cntr* (n-w=3). Statistics, Kruskal-Wallis test, p=0.5403 ns. Dunn's multiple comparison test, *Cntr* vs *ΔdIlp*s: p>0.9999 ns. **D.** *Graphs from top to bottom.* $AKT^{85\text{-}Ser505}$: $\Delta dIlp^{2}$ (dark blue, n-w=2, median=108.2), $\Delta dIlp^{3}$ (blue, n-w=1, median=111.8), $\Delta dIlp^{5}$ (dark purple, n-w=2, median=117.0), $\Delta dIlp^{7}$ (pink, n-w=2, median=115.8), $\Delta dIlp^{2\text{-}3}$ (light blue, n-w=2, median=115.1), $\Delta dIlp^{5,7}$ (light purple, n-w=1, median=119.5), $\Delta dIlp^{2\text{-}3,7}$ (green, n-w=2, median=118.2), *Cntr* (n-w=2). Statistics, Kruskal-Wallis test, p=0.3758 ns. Dunn's multiple comparison test, *Cntr* vs $\Delta dIlp^{2}$, $\Delta dIlp^{3}$: p>0.9999 ns; *Cntr* vs $\Delta dIlp^{5}$: p= 0.6573 ns; *Cntr* vs $\Delta dIlp^{7}$: p=0.6573 ns; *Cntr* vs $\Delta dIlp^{2\text{-}3}$: p=0.5088 ns; *Cntr* vs $\Delta dIlp^{5,7}$: p=0.2814 ns; *Cntr* vs $\Delta dIlp^{2\text{-}3,7}$: p=0.2188 ns. $AKT^{85\text{-}Thr342}$: $\Delta dIlp^{2}$ (dark blue, n-w=2, median=90.4), $\Delta dIlp^{3}$ (blue, n-w=1, median=90.3), $\Delta dIlp^{5}$ (dark purple, n-w=2, median=92.6), $\Delta dIlp^{7}$ (pink, n-w=2, median=93.9), $\Delta dIlp^{2\text{-}3}$ (light blue, n-w=2, median=94.7), $\Delta dIlp^{5,7}$ (light purple, n-w=1, median=87.5), $\Delta dIlp^{2\text{-}3,7}$ (green, n-w=2, median=102.4), *Cntr* (n-w=2). Statistics, Kruskal-Wallis test, p=0.2392 ns. Dunn's multiple comparison test, *Cntr* vs $\Delta dIlp^{2}$, $\Delta dIlp^{3}$, $\Delta dIlp^{5}$, $\Delta dIlp^{7}$, $\Delta dIlp^{2\text{-}3}$, $\Delta dIlp^{2\text{-}3,7}$: p>0.9999 ns; *Cntr* vs $\Delta dIlp^{5,7}$: p=0.4439 ns. P-MAPK: $\Delta dIlp^{2}$ (dark blue, n-w=2, median=119.9), $\Delta dIlp^{3}$ (blue, n-w=2, median=121.1), $\Delta dIlp^{5}$ (dark purple, n-w=2, median=133.9), $\Delta dIlp^{7}$ (pink, n-w=2, median=140.5), $\Delta dIlp^{2\text{-}3}$ (light blue, n-w=2, median=133.5), $\Delta dIlp^{5,7}$ (light purple, n-w=2, median=129.4), $\Delta dIlp^{2\text{-}3,7}$ (green, n-w=2, median=122.9), *Cntr* (n-w=2). Statistics, Kruskal-Wallis test, p=0.3939 ns. Dunn's multiple comparison test, *Cntr* vs $\Delta dIlp^{2}$, $\Delta dIlp^{3}$, $\Delta dIlp^{2\text{-}3,7}$: p>0.9999 ns; *Cntr* vs $\Delta dIlp^{5}$: p=0.4097 ns; *Cntr* vs $\Delta dIlp^{7}$: p=0.1095 ns; *Cntr* vs $\Delta dIlp^{2\text{-}3}$: p=0.4097 ns; *Cntr* vs $\Delta dIlp^{5,7}$: p=0.5179 ns. **E.** *Graphs from top to bottom.* $AKT^{85\text{-}Thr342}$: $\Delta dIlp^{2}$ (dark blue, n-w=3, median=117.5), $\Delta dIlp^{3}$ (blue, n-w=3, median=117.9), $\Delta dIlp^{5}$ (dark purple, n-w=3, median=104.4), $\Delta dIlp^{7}$ (pink, n-w=3, median=100.0), $\Delta dIlp^{2,7}$ (blue, n-w=3, median=98.1), $\Delta dIlp^{3,7}$ (green-blue n-w=3, median=96.8), $\Delta dIlp^{2\text{-}3}$ (light blue, n-w=3, median=96.1), $\Delta dIlp^{5,7}$ (light purple, n-w=3, median=91.5), *Cntr* (n-w=3). Statistics, Kruskal-Wallis test, p=0.5375 ns. Dunn's multiple comparison test, *Cntr* vs *ΔdIlp*s: p>0.9999 ns. P-MAPK: $\Delta dIlp^{2}$ (dark blue, n-w=3, median=74.6), $\Delta dIlp^{3}$ (blue, n-w=5, median=61.5), $\Delta dIlp^{5}$ (dark purple, n-w=3, median=75.9), $\Delta dIlp^{7}$ (pink, n-w=3, median=68.6), $\Delta dIlp^{2,7}$ (blue, n-w=3, median=86.7), $\Delta dIlp^{3,7}$ (green-blue, n-w=3, median=72.2), $\Delta dIlp^{2\text{-}3}$ (light blue, n-w=3, median=80.3), $\Delta dIlp^{5,7}$ (light purple, n-w=3, median=78.1), *Cntr* (n-w=3). Statistics, Kruskal-Wallis test, p=0.0534 ns. Dunn's multiple comparison test, *Cntr* vs $\Delta dIlp^{2}$: p=0.1432 ns; *Cntr* vs $\Delta dIlp^{3}$: p=0.0104 *; *Cntr* vs $\Delta dIlp^{5}$: p=0.5110 ns; *Cntr* vs $\Delta dIlp^{7}$: p=0.0595 ns; *Cntr* vs $\Delta dIlp^{2,7}$, $\Delta dIlp^{5,7}$ p>0.9999 ns; *Cntr* vs $\Delta dIlp^{3,7}$: p=0.1079 ns; *Cntr* vs $\Delta dIlp^{2\text{-}3}$: p=0.6061 ns.

**(F).** The weight (in mg per adult) was calculated from n adult pools (n-p). Each pool contained $n_{adult}$ ≥9. *Cntr* (grey, $N_{adults}$=325, n-p=7, median=1.13), $\Delta dIlp^{2}$ (dark blue, $N_{adults}$=350, n-p=9, median=0.85), $\Delta dIlp^{3}$ (blue, $N_{adults}$=212, n-p=5, median=0.82), $\Delta dIlp^{5}$ (purple, $N_{adults}$=272, n-p=6, median=0.97), $\Delta dIlp^{7}$ (pink, $N_{adults}$=325, n-p=7, median=0.87), $\Delta dIlp^{2\text{-}3}$ (light blue, $N_{adults}$=407, n-p=13, median=0.93), $\Delta dIlp^{2,7}$ (blue, $N_{adults}$=407, n-p=13, median=1.04), $\Delta dIlp^{3,7}$ (green-blue, $N_{adults}$=332, n-p=7, median=0.94), $\Delta dIlp^{5,7}$ (light purple, $N_{adults}$=272, n-p=6, median=0.98), $\Delta dIlp^{2\text{-}3,7}$ (green, $N_{adults}$=391, n-p=12, median=0.78). Statistics, Kruskal-Wallis test, p<0.0001 ****. Dunn's multiple comparison test, *Cntr vs* $\Delta dIlp^{2}$: p=0.0014 **; *Cntr vs* $\Delta dIlp^{3}$: p=0.0015 **; *Cntr vs* $\Delta dIlp^{5}$, $\Delta dIlp^{2,7}$, $\Delta dIlp^{5,7}$: p>0.9999 ns; *Cntr vs* $\Delta dIlp^{7}$: p=0.0007 ***; *Cntr vs* $\Delta dIlp^{2\text{-}3}$: p=0.0304 *; *Cntr vs* $\Delta dIlp^{3,7}$: p=0.7729 ns; *Cntr vs* $\Delta dIlp^{2\text{-}3,7}$: p<0.0001 ****.

**(G).** Here are plotted the protein levels in adult haemolymph (in arbitrary units, A.U.). The adults were reared on YF at 20°C or 28°C. For each genotype $N_{exp}$=3. *Cntr* (grey, $median^{20°C}$=0.059, $median^{28°C}$=0.048), $\Delta dIlp^{2}$ (dark blue, $median^{20°C}$=0.028, $median^{28°C}$=0.011), $\Delta dIlp^{3}$ (blue, $median^{20°C}$=0.035, $median^{28°C}$=0.011), $\Delta dIlp^{5}$ (purple, $median^{20°C}$=0.004, $median^{28°C}$=0.033), $\Delta dIlp^{7}$ (pink, $median^{20°C}$=0.017, $median^{28°C}$=0.025), $\Delta dIlp^{2\text{-}3}$ (light blue, $median^{20°C}$=0.007, $median^{28°C}$=0.030), $\Delta dIlp^{2,7}$ (blue, $median^{20°C}$=0.034, $median^{28°C}$=0.020), $\Delta dIlp^{3,7}$ (green-blue, $median^{20°C}$=0.029, $median^{28°C}$=0.016), $\Delta dIlp^{5,7}$ (light purple, $median^{20°C}$=0.018, $median^{28°C}$=0.032), $\Delta dIlp^{2\text{-}3,7}$ (green, $median^{20°C}$=0.039, $median^{28°C}$=0.042). Statistics, Kruskal-Wallis test, p=0.1727 ns. Dunn's multiple comparison test, *Cntr vs* $\Delta dIlp^{2}$: $p^{20°C}$=0.4631 ns, $p^{28°C}$=0.7020 ns; *Cntr vs* $\Delta dIlp^{3}$: $p^{20°C}$>0.9999 ns, $p^{28°C}$=0.5151 ns; *Cntr vs* $\Delta dIlp^{5}$: $p^{20°C}$=0.1837 ns, $p^{28°C}$>0.9999 ns; *Cntr vs* $\Delta dIlp^{7}$, $\Delta dIlp^{2,7}$, $\Delta dIlp^{3,7}$, $\Delta dIlp^{2\text{-}3,7}$: $p^{20°}$, $p^{28°C}$>0.9999 ns; *Cntr vs* $\Delta dIlp^{2\text{-}3}$: $p^{20°C}$=0.0270 *, $p^{28°C}$>0.9999 ns.

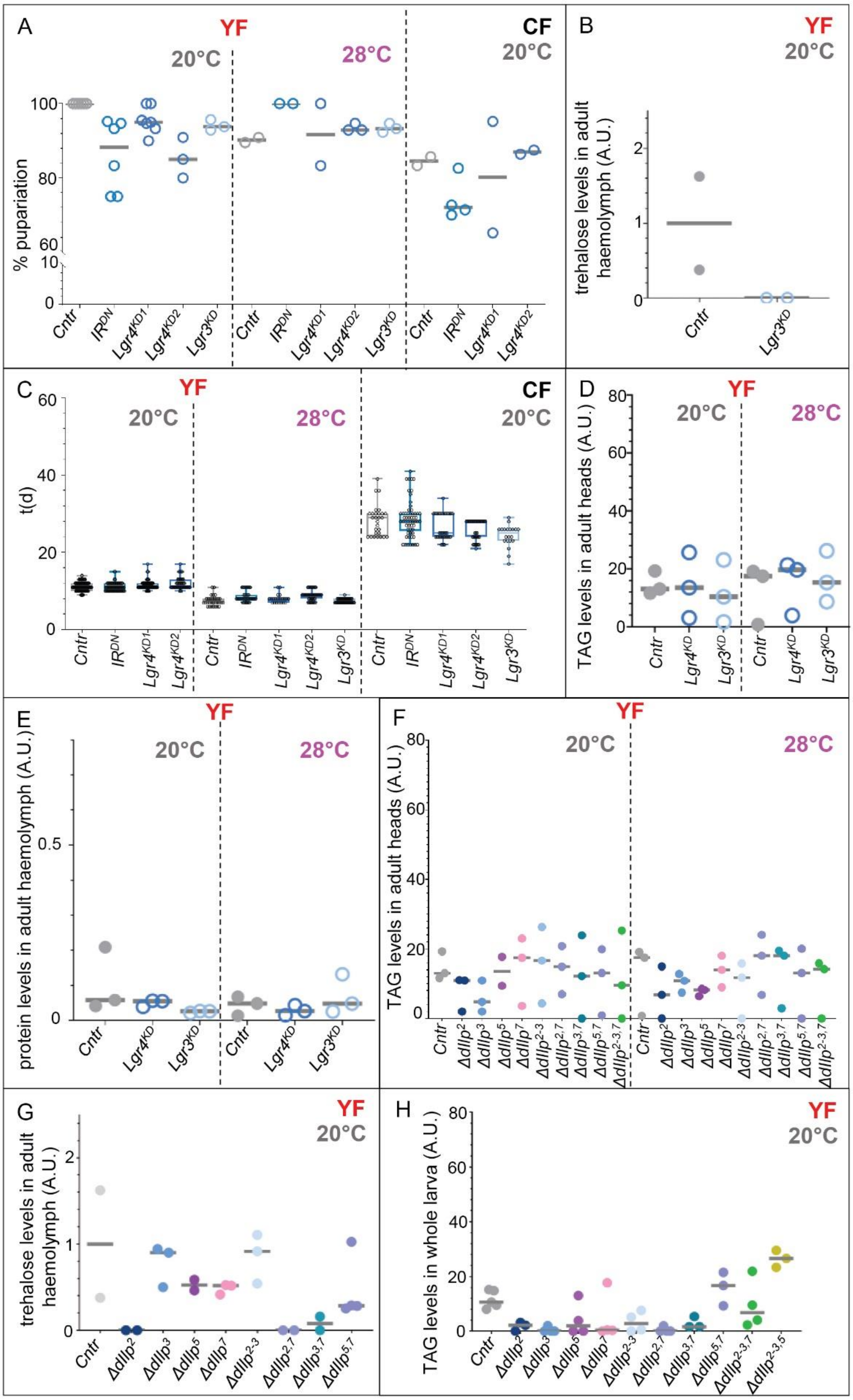


**Supplementary Figure S4. Screen of the predicted candidate dIlp7 receptors**

**(A).** Plotted are the pupariation rate (%) of larvae reared on YF (red) at 20°C or 28°C, or on CF (black, right) at 20°C. The genotypes shown are *dIlp2-Gal4/+* (*Control*, *Cntr*, grey circles), *dIlp2>>IR*$^{DN}$ (*IR*$^{DN}$, blue circles), *dIlp2>>Lgr4RNAi (KK102681)* (*Lgr4*$^{KD1}$, bright-blue circles), *dIlp2>>Lgr4RNAi*

*(KK108915)* (*Lgr4*$^{KD2}$, bright-blue circles) and *dIlp2>>Lgr3RNAi* (*Lgr3*$^{KD}$, , light-blue circles). For the *Lgr3*$^{KD}$ data on CF, see Fig.4B. One dot= one experiment; the median is depicted as a grey bar. *Cntr* (YF, median$^{20°C}$=100%, median$^{28°C}$=90.2%; CF, median$^{20°C}$=84.5%), *IR*$^{DN}$ (YF, median$^{20°C}$=88.3%, median$^{28°C}$=100%; CF, median$^{20°C}$=71.4%), *Lgr4*$^{KD1}$ (YF, median$^{20°C}$=95.0%, median$^{28°C}$=91.7%; CF, median$^{20°C}$=80.2%), *Lgr4*$^{KD2}$ (YF, median$^{20°C}$=85.0%, median$^{28°C}$=92.9%; CF, median$^{20°C}$=86.9%), *Lgr3*$^{KD}$ (YF, median$^{20°C}$=93.8%, median$^{28°C}$=93.3%; CF, median$^{20°C}$=62.5%). Each experiment included $n_{larvae}$ ≥10 (20°C) or $n_{larvae}$ ≥6 (28°C). Statistics, Kruskal-Wallis, p=0.0025 **. Dunn's multiple comparison test, *Cntr* vs *IR*$^{DN}$: p$^{20°C}$=0.0028 **, p$^{28°C}$=0.0688 ns (YF), p$^{20°C}$=0.7577 ns (CF); *Cntr* vs *Lgr4*$^{KD1}$: p$^{20°C}$=0.3159 ns, p$^{28°C}$>0.9999 ns (YF), p$^{20°C}$>0.9999 ns (CF); *Cntr* vs *Lgr4*$^{KD2}$: p$^{20°C}$=0.0050 **, p$^{28°C}$=0.8774 ns (YF), p$^{20°C}$ >0.9999 ns (CF); *Cntr* vs *Lgr3*$^{KD}$: p$^{20°C}$=0.2534 ns, p$^{28°C}$=0.8774 ns (YF).

**(B).** Trehalose levels (in arbitrary units, A.U.) have been measured in *FOXO*$^{mCherry}$ (*Control*, *Cntr*, grey) and *dIlp2>>Lgr3RNAi* (*Lgr3*$^{KD}$, light-blue circles) adult haemolymph. Adults were fed on YF at 20°C. The levels of trehalose have been normalised to *Cntr* ($N_{exp}$=2, median=1). *Lgr3*$^{KD}$ ($N_{exp}$=2, median=0). Statistics, Mann-Whitney test, p=0.3333 ns.

**(C).** Plotted is the developmental speed (in days, d) of larvae reared on YF (red, left) at 20°C or 28°C, or on CF (black, right) at 20°C. The genotypes shown are *dIlp2-Gal4/+* (*Control*, *Cntr*, grey), *dIlp2>>IR*$^{DN}$ (*IR*$^{DN}$, blue circles), *dIlp2>>Lgr4RNAi (KK102681)* (*Lgr4*$^{KD1}$, bright-blue circles), *dIlp2>>Lgr4RNAi (KK108915)* (*Lgr4*$^{KD2}$, bright-blue circles) and *dIlp2>>Lgr3RNAi* (*Lgr3*$^{KD}$, light-blue circles). For the *Lgr3*$^{KD}$ data on YF at 20°C, see Fig.4A. One dot= one experiment; the median is depicted as a grey bar. *Cntr* (YF, median$^{20°C}$=11d, median$^{28°C}$=7d; CF, median$^{20°C}$=29d), *IR*$^{DN}$ (YF, median$^{20°C}$=11d, median$^{28°C}$=8d; CF, median$^{20°C}$=28d), *Lgr4*$^{KD1}$ (YF, median$^{20°C}$=11d, median$^{28°C}$=8d; CF, median$^{20°C}$=25d), *Lgr4*$^{KD2}$ (YF, median$^{20°C}$=11d, median$^{28°C}$=9d; CF, median$^{20°C}$=28d), *Lgr3*$^{KD}$ (YF, median$^{28°C}$=7d; CF, median$^{20°C}$=25d). Each experiment included $n_{larvae}$ ≥10 (20°C) or $n_{larvae}$ ≥6 (28°C). Statistics, p<0.0001 ****. Dunn's multiple comparison test, *Cntr vs IR*$^{DN}$: p$^{20°C}$=0.7297 ns, p$^{28°C}$=0.0006 *** (YF), p$^{20°C}$>0.9999 ns (CF); *Cntr vs Lgr4*$^{KD1}$: p$^{20°C}$=0.2012 ns, p$^{28°C}$=0.2994 ns (YF), p$^{20°C}$=0.6226 ns (CF); *Cntr vs Lgr4*$^{KD2}$: p$^{20°C}$=0.0121 *, p$^{28°C}$<0.0001 **** (YF), p$^{20°C}$=0.0162 * (CF); *Cntr vs Lgr3*$^{KD}$: p$^{28°C}$>0.9999 ns (YF), p$^{20°C}$=0.0016 ** (CF).

**(D).** Triacylglycerid (TAG) levels (in arbitrary units, A.U.) have been measured in *FOXO*$^{mCherry}$ (*Control*, *Cntr*, grey), *dIlp2>>Lgr4RNAi* (KK108915) (*Lgr4*$^{KD}$, light-blue circles) and *dIlp2>>Lgr3RNAi* (*Lgr3*$^{KD}$, light-blue circles) adult heads. Adults were fed on YF at 20°C or 28°C. For each genotype, $N_{exp}$=3. *Cntr* (median$^{20°C}$=13.04, median$^{28°C}$=17.51), *Lgr3*$^{KD}$ (median$^{20°C}$=10.38, median$^{28°C}$=15.30). Statistics, Kruskal-Wallis test, p=0.9663 ns. Dunn's multiple comparison test, *Cntr vs Lgr4*$^{KD}$, *Lgr3*$^{KD}$: p>0.9999 ns.

**(E).** Trehalose levels (in arbitrary units, A.U.) have been measured in *FOXO*$^{mCherry}$ (*Control*, *Cntr*, grey) and *ΔdIlp* mutant adult haemolymph. Adults were fed on YF at 20°C. The levels of trehalose have been normalised to *Cntr* ($N_{exp}$=2, median=1). *ΔdIlp*$^{2}$ (dark blue, $N_{exp}$=2, median=0), *ΔdIlp*$^{3}$ (blue, $N_{exp}$=3, median=0.903), *ΔdIlp*$^{5}$ (dark purple, $N_{exp}$=2, median=0.525), *ΔdIlp*$^{7}$ (pink, $N_{exp}$=3, median=0.519), *ΔdIlp*$^{2-3}$ (light blue, $N_{exp}$=3, median=0.917), *ΔdIlp*$^{2,7}$ (blue, $N_{exp}$=2, median=0), *ΔdIlp*$^{3,7}$ (green-blue, $N_{exp}$=2, median=0.079), *ΔdIlp*$^{5,7}$ (light purple, $N_{exp}$=4, median=0.286). Statistics, Kruskal-Wallis test, p=0.0455 *. Dunn's multiple comparison test, *Cntr vs ΔdIlp*$^{2}$: p=0.3636 ns; *Cntr vs ΔdIlp*$^{3}$, *ΔdIlp*$^{5}$, *ΔdIlp*$^{7}$, *ΔdIlp*$^{2-3}$, *ΔdIlp*$^{5,7}$: p>0.9999 ns.

**(F).** Triacylglycerid (TAG) levels (in arbitrary units, A.U.) have been measured in *FOXO*$^{mCherry}$ (*Control*, *Cntr*, grey) and *ΔdIlp* mutant larva haemolymph. Larvae were fed on YF at 20°C. *Cntr* ($N_{exp}$=5, median=10.58), *ΔdIlp*$^{2}$ (dark blue, $N_{exp}$=3, median=2.15), *ΔdIlp*$^{3}$ (blue, $N_{exp}$=4, median=0.00), *ΔdIlp*$^{5}$ (dark purple, $N_{exp}$=4, median=1.91), *ΔdIlp*$^{7}$ (pink, $N_{exp}$=4, median=0.37), *ΔdIlp*$^{2-3}$ (light blue, $N_{exp}$=4, median=2.80), *ΔdIlp*$^{2,7}$ (blue, $N_{exp}$=4, median=0.00), *ΔdIlp*$^{3,7}$ (green blue, $N_{exp}$=3, median=1.61), *ΔdIlp*$^{5,7}$ (light purple, $N_{exp}$=3, median=16.65), *ΔdIlp*$^{2-3,7}$ (green, $N_{exp}$=4, median=6.77), *ΔdIlp*$^{2-3,5}$ (yellow, $N_{exp}$=3, median=26.57). Statistics, Kruskal-Wallis test, p=0.0034 **. Dunn's multiple comparison test, *Cntr vs ΔdIlp*$^{2}$: p=0.7057 ns; *Cntr vs ΔdIlp*$^{3}$: p=0.0570 ns; *Cntr vs ΔdIlp*$^{5}$: p=0.6846 ns; *Cntr vs ΔdIlp*$^{7}$: p=0.7337 ns; *Cntr vs ΔdIlp*$^{2-3}$: p=0.8983 ns; *Cntr vs ΔdIlp*$^{2,7}$: p=0.0518 ns; *Cntr vs ΔdIlp*$^{3,7}$, *ΔdIlp*$^{5,7}$, *ΔdIlp*$^{2-3,7}$, *ΔdIlp*$^{2-3,5}$: p>0.9999 ns.

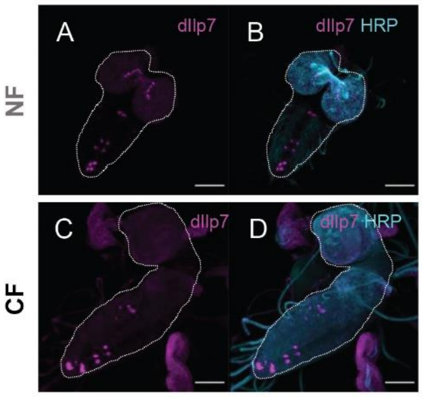


**Supplementary Figure S5. D7Ns accumulate dIlp7 peptide on CF**
Representative Z-stack confocal images (A, B, stack= 100μm; C, D, stack=66.96μm) of third-instar larva CNS probed for dIlp7 (magenta) and HRP (cyan; B, D). Larvae were reared on NF (A, B) or CF (C, D). Scale bars, 50μm.
